# Differential expression of spatiotemporal sleep spindle clusters in ageing

**DOI:** 10.1101/2025.05.19.654222

**Authors:** Liisa Raud, Martijn J. B. Smits, Markus H. Sneve, Hedda T. Ness, Line Folvik, Björn Rasch, Anders M. Fjell

## Abstract

**Objectives:** Sleep spindles are potential biomarkers for memory decline in aging. However, significant within-person variability in spindle attributes complicates their utility in predicting cognitive deterioration. This study aimed to uncover distinct spindle types and their relevance to memory decline using data-driven clustering.

**Methods:** Polysomnography was collected from younger (n = 43, ages 20-45 years) and older cognitively healthy adults (n = 34, ages 60-81 years). Spindles were clustered into four groups using multiple features and spatiotemporal context, irrespective of participant age.

**Results:** Resulting clusters were hierarchically defined by the sleep stage, slow oscillation concurrence, and hemisphere. Stage N3 spindles (15%; predominantly coinciding with slow oscillations) formed a distinct group, followed by N2 spindles coinciding with slow oscillations (27%). Remaining N2 spindles were categorized into unilateral (41%) and bilateral clusters (17%). In older adults, there was a reduced proportion of N2 bilateral spindles and an increased proportion of N2 spindles concurrent with slow oscillations. Reduced proportion of N2 bilateral spindles was associated with better composite memory performance in younger adults, whereas higher spindle power, regardless of cluster belonging, was associated with reduced memory benefit from sleep compared with wakefulness.

**Conclusions:** Our results indicate differing expression of spatiotemporal spindle clusters in older age, as well as intertwined dynamics between spindle propagation, SO concurrence, and frequency shifts in ageing. In addition, spindle heterogeneity aligned with global sleep stage dynamics. These results emphasize the interconnectedness of spindle activity with overall sleep patterns, underscoring the importance of spatiotemporal context within and across sleep stages.

**Statement of significance:** This study used data-driven clustering to explore sleep spindles as potential markers for age-related memory decline. We identified spindle clusters determined by sleep stage, slow oscillation concurrence, and hemisphere propagation. Notably, older adults showed altered expression of these clusters, indicating age-specific dynamics. Further research should focus on distinguishing spindle deterioration from broader sleep changes in older age. Such insights could pave the way for early detection and intervention strategies in cognitive decline, highlighting sleep’s crucial role in maintaining cognitive health and resilience in aging populations. These findings hold promise for developing targeted approaches to enhance mental wellness and quality of life in older adults.

## INTRODUCTION

As people age, changes in sleep architecture become prominent. These include alterations in the duration and quality of various sleep stages, as well as in specific physiological features such as sleep spindles and slow oscillations^1–3^. At the same time, aging is often accompanied by a decline in memory capabilities, significantly impacting daily life and overall well-being^4,5^. It has been suggested that sleep plays a crucial role in memory consolidation across the human lifespan^6–8^. Consequently, establishing a link between quantifiable sleep markers and memory could be promising for the early detection of cognitive decline in the older adults. Sleep spindles, in particular, have emerged as promising candidates for predicting memory decline in both healthy and pathological aging^9–17^ . However, the ability to predict memory changes across individuals may be compromised by the significant within-individual heterogeneity in spindle attributes throughout the night^18^.

Sleep spindles are distinctive bursts of oscillatory brain activity that can be recorded by electroencephalography (EEG)^19^. These events occur during non-rapid eye movement (NREM) sleep and are characterized by a frequency range of approximately 9-16 Hz and a duration of about 0.5-2 seconds. The rapid changes in cellular Ca2+ concentrations triggered by spindle events make them relevant for various types of neural reorganization^19,20^, including spike-time dependent plasticity^21^, long-term potentiation and depression^22–24^, and dendritic spine formation^25^. Sleep spindles also play a significant role in active systems consolidation theory, where, alongside concurrent slow oscillations (SOs) and hippocampal ripples, it is speculated that they facilitate communication between the hippocampus and the neocortex, essential for the transfer and long-term storage of new information^7,26,27^.

Compared to young adults, the older adults have reduced spindle density, power, duration, and dominant frequency^19,28–33^. While age may influence the spindle-memory association in healthy individuals^34,35^, deficiencies in spindle expression could precede age-related brain changes as deterioration progresses^10^. To identify healthy adults at risk for dementia, Orlando et al. (2023)^36^ utilized a clustering approach on averaged spindle attributes and found a group of individuals with low spindle density, duration, and power, who had the greatest memory decline. This approach could be enhanced by considering the extensive spindle heterogeneity within individuals. In our study, we adopted a similar clustering approach but focused on clustering at the level of individual spindles to capture their variability within each person.

The sources of spindle variability throughout the night are manifold. The most commonly considered attribute is spindle frequency, distinguishing between slow (9-12 Hz) and fast (12-16 Hz) spindles^28,37,38^. There is ongoing debate about whether these spindles represent functionally distinct types^18^. Generally, stronger associations are found between memory and fast spindles^39^, although a recent large-scale study suggests that both spindle types may predict cognitive performance in older adults^40^.

A related approach differentiates spindles based on their scalp topography, distinguishing between frontal and parietal spindles^37^. However, findings from larger electrode coverages and intracranial recordings indicate more variable spatial organization^18,41,42^, suggesting that a coarse distinction between local and global spindles might also be appropriate.

The temporal context of spindles is another significant factor. Spindles occurring on the up-phase of SOs are particularly relevant for memory and appear to deteriorate with age^43,44^. This presents a paradox, as SOs are the hallmark of stage N3 sleep, while spindles are most prominent in N2 sleep^18^, making their co-occurrence, as measured by EEG, relatively rare^41^. The timing of spindle events during the night is also pertinent^45^, as spindle power varies non-linearly throughout the night^46,47^. Additionally, the timing within sleep cycles is relevant^48,49^.

Furthermore, spindle characteristics may be influenced by the type of thalamocortical projections; core projections are limited to deeper cortical layers, while matrix projections reach the upper layers and are more likely captured by EEG electrodes^50,51^.

Lastly, the information in the post-spindle signal might be as critical as the spindle attributes. Given that spindles are highly synchronous oscillatory events, they may have limited information-processing capacity^52^ and may rather serve as gating phenomena^53–56^ that initiate subsequent information transfer in the cortex^57^. This is indirectly evidenced by the so-called spindle refractory period, during which incoming information seems not to be processed within a few seconds after a spindle event, as if blocked by ongoing computations^58^.

In sum, there are significant sources of variability in spindle expression throughout the night within each person. Notably, this variability is not random, as specific features tend to co-occur^18^. For example, slow spindles are more localized to the frontal areas, while faster spindles are more prominent in the temporo-parietal regions^37^. At the same time, stage N2 spindles generally exhibit higher power^41^ or faster frequency^18^ compared to those in N3. Therefore, data-driven clustering of single spindles based on an array of different attributes may uncover functionally distinct spindle types.

On the other side of the equation, it remains unclear which aspects of age-related memory decline are most influenced by changes in sleep. Whereas sleep spindles may be preferably associated with procedural memory^15^, ageing appears to affect more profoundly sleep based consolidation of declarative memory^59^. In addition, several moderating factors have been identified, including older age, where the beneficial effects of sleep on memory performance diminish or even disappear^60^. Studies on long-term declarative memory suggest that aging also affects the durability of already learned associations^61–63^. However, the role of sleep on the longevity of episodic memories in older adults, and how these sleep-memory interactions influence daily functioning, remains an open question.

In this study, we leveraged single spindle activity and various memory tasks to identify spindle types relevant to age-related memory decline in healthy participants. Specifically, we used multiple episodic memory tasks, combining specific memory gain in sleep over wakefulness with delayed recall assessments and subjective memory problems. While our analysis was data-driven and exploratory, it was guided by three core hypotheses. First, we anticipated heterogeneity in spindle characteristics, which would allow them to be grouped into distinct types. Second, we expected age differences in the expression of these spindle types. Third, we expected that parsing this spindle heterogeneity into distinct clusters may help to pinpoint which spindle types are the best predictors of age-related memory decline as composite score and for sleep-associated memory consolidation.

## METHODS

### Sample

Data was analyzed from 77 healthy adults, divided into younger (n = 43; 20-45 years, mean age = 27, sd = 5, 23/20 males/females) and older (n = 34; 60-81 years, mean age = 68, sd = 5, 20/14 males/females) age groups. Initially, 92 individuals were enrolled in the study, of which 82 had PSG data available. From these, data was discarded due to bad EEG quality (n = 2) and high depression scores (n = 3; Becks Depression Inventory >= 16 or Geriatric Depression Scale >= 9). All participants were fluent in Norwegian, right-handed, with normal or corrected vision and no history of severe psychiatric or neurological disorders, traumatic or enhanced brain injury, sleep disorders, and no current use of medications known to affect the nervous system. All participants signed an informed consent approved by the Regional Ethical Committee of South Norway (REK 2010/3407). The main recruiting strategies included targeted Social Media advertisement, flyers, and posters at selected places (e.g. senior centers). Participants received monetary compensation for the participation.

### General procedure

This study is part of a larger project investigating memory consolidation processes at different timescales and their possible relation to memory decline in aging. The project design included two experimental sessions per participant, divided into an awake and a sleep condition (Figure 1). Each session consisted of several elements occurring in this order: memory encoding task, eight-alternative forced choice retrieval task (AFC-1), 12 hours waiting period, memory retrieval task, AFC-2, and online AFC-3 around six days later. The experimental encoding and retrieval tasks took place within MRI scanner, with results reported elsewhere^62–65^.

**Figure 1.**
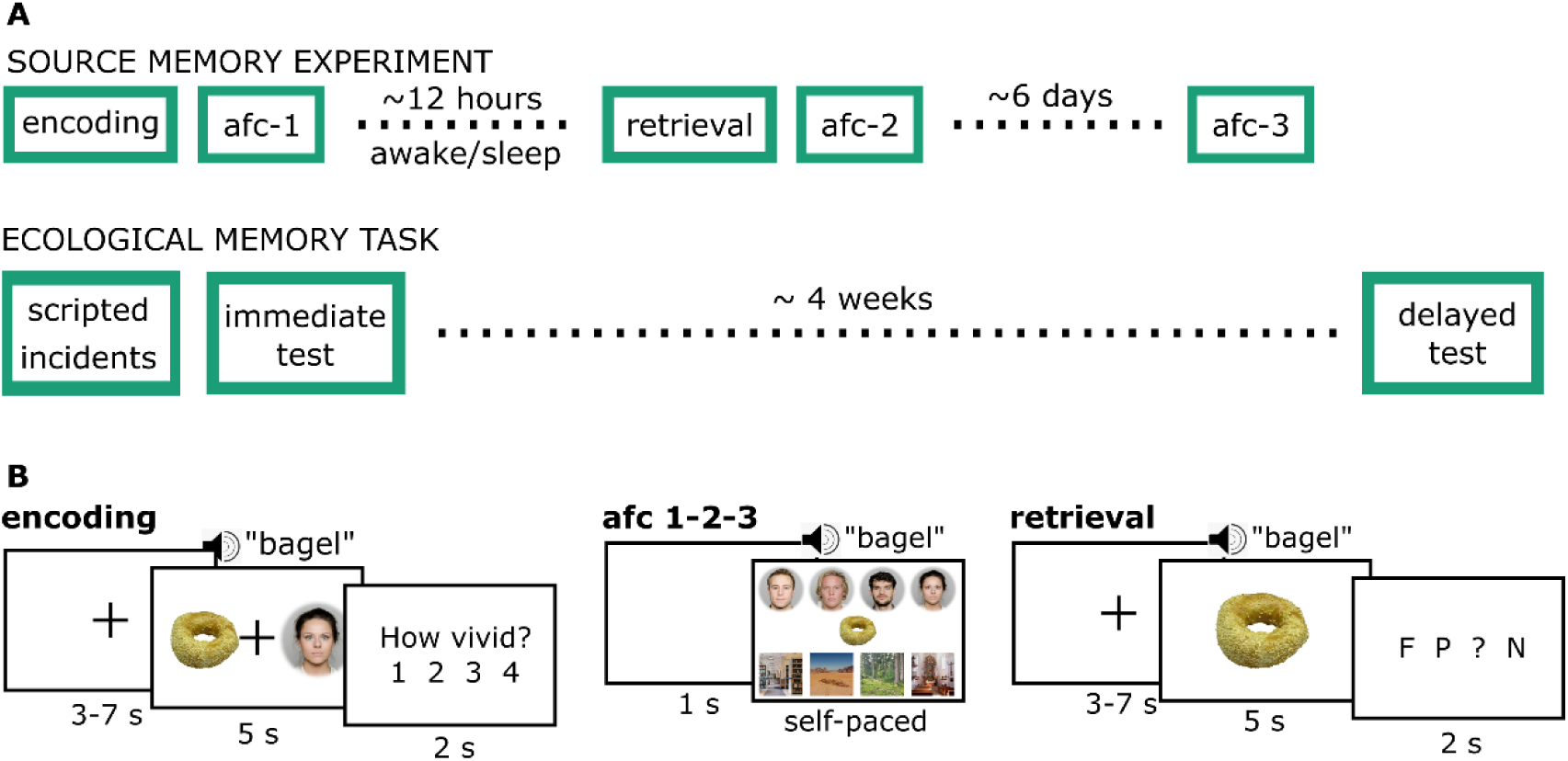
(A) Timelines of the experimental ecological and source memory tasks. The source memory experiment was admitted in two conditions, once with awake 12-hour period during the day and once during the night. In the latter condition, participants wore ambulatory polysomnography device to record sleep-EEG. The scripted part of the ecological memory task and the immediate testing were incorporated in a separate session for neuropsychological testing. (B) Experimental source memory task phases. During encoding, an item was represented with a face or a place (visually and auditorily) and participants were asked to form a mental association between the two. The vividness of the association was rated after each trial. In AFC, participants were shown each item and made to choose which face/place it was associated with. During the retrieval phase, the participants received audio-visual presentation of each item and were asked whether it had been previously associated with a face (F), place (P), they did not remember (?), or the item is new (N). AFC – (eight-)alternative forced choice task. The face-stimuli were drawn from the Oslo Face database^66,67^, the item-stimuli were drawn from the Bank of Standardized stimuli^68,69^, and the place-stimuli from Pixabay and Google searches, marked for non-commercial use with changes. The face images presented here are not identical to those used in the task but includes a selection of images in which the individuals have consented to their faces being publicly displayed.

The order of the experimental sessions (wake vs. sleep) was counterbalanced across participants. During the sleep condition, participants arrived at the hospital either 7 or 9 p.m. and subsequently began the memory encoding task at approximately 8 or 10 p.m., respectively. After completing the evening memory tasks in the MRI scanner, they were equipped with a polysomnography (PSG) device and directed to sleep in the hotel next to the hospital. They returned next morning at 8 or 10 a.m. to complete the memory retrieval tasks. Prior to the experimental night, the participants had a habituation night with the PSG at the same hotel.

In addition to the experimental sessions, participants filled online questionnaires and visited the lab for neuropsychological testing. Relevant for this study, they filled in Becks Depression Inventory, Geriatric Depression Inventory, Pittsburgh Sleep Quality Index, and Everyday Memory Questionnaire. A custom ecological memory test procedure was integrated into this testing session, probing participants at the end of the first session and four weeks later by phone.

### Memory evaluation

Six different memory scores were extracted from the different memory evaluation procedures, which were later combined using principal component analysis (PCA).

*Source memory experiment.* The source memory experiment contributed with three conceptually different scores, named “source recall”, “delayed source recall”, and “sleep gain”. The details of the source memory task (Figure 1B) are described previously^62–65^. Shortly, during the encoding phase, participants were shown pictures of 128 items, each together with a specific face or place (4 faces and 4 places in total). They were instructed to imagine an association between the item and the face/place, and then to evaluate the vividness of this association on the scale 1-4. Directly after the encoding task, their learning performance was tested with a forced choice task (AFC), in which the participant had to indicate, for each item, which face or place it had been paired with. After 12 hours, participants returned for a retrieval test, in which they were shown each item and several new items, and they had to indicate whether it had been previously associated with a face, place, they did not remember, or the item was new. The experimental memory score (“source recall”) was calculated by counting the correctly remembered face/place associations, subtracting the number of incorrect associations, divided by the total number of items, and lastly, averaged over the wake and sleep sessions for more robust estimation. This score thus reflects the percentage of correctly remembered associations after 12 hours, corrected for guessing. Given that the retrieval task prompts for the context in which each item association was formed, it is putatively reflecting episodic source memory^70^.

The second memory score (“dealyed source recall”) was also derived from the experimental procedure, particularly the 8-alternative forced choice tasks (AFC-1-2-3). In this task, participants were shown each item together with all four faces and four places. They had to choose, which face or place the item had been associated with during the encoding phase. This test was administered three times: first directly after encoding, then after the 12-hour period (directly after the in-scanner retrieval task described previously), and lastly, after about six days using an online version of the task. The memory score was calculated as the percentage of accurate associations in all three timepoints (hit-hit-hit), thus indicating the percentage of associations correctly remembered after six days, putatively reflecting the ability to form durable memories^62,63^.

The third memory score, “sleep gain,” represents the benefit of a 12-hour sleep period compared to the same duration of wakefulness and is calculated as the difference in source recall accuracy (corrected for guessing) between sleep and wake conditions. A positive value indicates improved memory performance after sleep relative to the awake period, serving as an index of sleep-driven memory consolidation. This retrieval test focused on category associations rather than specific faces and places, making it well-suited for assessing gist memory formation and consolidation, typically linked to sleep^71–73^. However, since this test was not administered immediately after learning, the sleep-wake difference might be influenced by baseline variations before the 12-hour period. To address this, we additionally calculated the sleep gain effect using AFC-1 (baseline) and AFC-2 test performances (Figure 1). First, AFC-2 performance was calculated as the percentage of items remembered at both time points, relative to baseline AFC-1 performance. Then, difference scores between sleep and wake conditions were calculated.

*Ecological memory task.* This task was a custom ecological memory test, designed to objectively measure everyday attentional and memory performance, and contributed with two memory scores: “everyday recall”, and “dealyed everyday recall”. This was implemented in the separate lab-visit, when the participants came in for neuropsychological testing (not used in this study). During this session, the experimenters highlighted ten previously scripted specific behaviors or details in the environment, for example asking the participant to hold a pen during a procedure, pointing out the location of the restrooms, commenting on the type of the music played during waiting, etc. At the end of the session, the participants were asked to recall what they did from the beginning of session until the start of the session, including as much detail as possible. After this spontaneous narration, participants were probed directly about specific episodes that they had omitted from their narratives. This procedure was repeated four weeks later via a phone call. With participants’ consent, their narratives were recorded and later transcribed by the experimenter following a 13-item questionnaire. The items were coded ‘correct’ if remembered correctly, ‘incorrect’ if participant indicated they had forgotten or gave a wrong answer, or ‘irrelevant’ if the answer was indefinite or too abstract for direct transcription, or when there was ambiguity whether this action was correctly performed during the session. All irrelevant items were discarded, and the percentage of correct responses to the remaining items served as the fourth (“everyday recall”) and fifth (“dealyed everyday recall”) memory score, reflecting immediate and delayed recall, respectively.

*Everyday Memory Questionnaire.* The last score (“emq”) was derived from the Everyday Memory Questionnaire^74,75^, which measures self-reported memory failures in everyday life. This 28-item questionnaire probes participants for the frequency of daily memory lapses, such as loosing items, forgetting appointments, etc. We calculated the sum of all items, and higher sum scores indicate worse perceived memory problems.

*Principal component analysis.* All six scores were combined into a single memory score using PCA in R (v. 4.2.1). The calculation was done by a singular value decomposition on the zero-centered and scaled (to have unit variance) data matrix. Prior to the PCA, missing values (n=28, constituting 6% across all memory scores) were imputed based on available values using k nearest neighbors method, implemented by the impute package^76^.

### PSG acquisition

PSG was recorded with two different devices: SomnoTouch Resp (SOMNOmedics AG, Germany), and Nox A1 (Resmed, USA). With SomnoTouch Resp (n = 64), data was collected over 4 EEG electrodes (C3, C4, M1, M2), 3 bipolar chin-EMG electrodes, and 2 EOG electrodes. Data was sampled at 256 Hz, and bandpass software filters for EEG (0.1-35 Hz) and EMG (0.1-100 Hz) were applied during data export. With Nox A1 device (n = 13), data was sampled at 500 Hz over 8 EEG electrodes (C3, C4, F3, F4, O1, O2, M1, M2), 3 chin-EMG electrodes and 2 EOG electrodes. Active electrodes were referenced to the opposing mastoids during data export.

### PSG preprocessing

PSG data was harmonized and preprocessed with MNE v.0.19.2^77^. Channels C3 and C4 were retained and referenced to M2 and M1, respectively, as these were common for both devices. The data was re-sampled to 200 Hz with automatic anti-aliasing filter. The Nox A1 data was additionally filtered between 0.1 and 35 Hz using non-pass zero-phase Hamming window bandpass filter (−6db cutoff at 0.05 and 39.38 Hz).

### Sleep staging

Continuous PSG was epoched into 30 second segments. For reproducible workflow, automatic sleep staging was implemented using the YASA toolbox^78^ (0.6.3) in Python 3.8, in which each segment was assigned a sleep stage based on the AASM guidelines^79^. Manually staged data is available, scored by a student lab member. The average overlap between automatic and human staged data was 77% (sd = 11), which rose to 80% (sd = 7) after discarding three outliers with <60% overlap, and is in the range of overlap between different human raters^80^.

### Event detection

Sleep spindles were also detected with the YASA toolbox. Spindle detection was limited to stage N2 and N3 and was applied to both C3 and C4 electrodes, with the frequency band of 9-16 Hz. The algorithm follows coarsely Lacourse et. al. 2018^81^, with major diversions explained in YASA online tutorial^82^. The raw signal was bandpass filtered between 1-30 Hz. Then, data was segmented into consecutive epochs of 2 seconds with 200 ms overlap, and submitted to short-term Fourier Transform to calculate the frequency power spectrum per epoch. We adopted two thresholds to detect potential spindle events. The first threshold was based on a sliding window (300 ms in steps of 100 ms) correlation between the sigma-filtered signal and the bandpass filtered EEG. Specifically, the Pearson correlation coefficient had to be >=0.65, ensuring that the detected spindles would also be visible in the raw EEG – the main criteria for manual detection based on visual inspection^81^. The second threshold was based on a sliding window root mean square (RMS) of the sigma-filtered EEG (300 ms in steps of 100 ms). This threshold was exceeded, if the RMS exceeded 1.5 standard deviations from the mean RMS, calculated without the 10% of highest and lowest values to reduce the bias due to potential artefacts. Note that the third threshold (implemented by default in YASA) that considers the relative power of the sigma frequency band with respect to the total broadband power was discarded in our implementation. This is because we intended the algorithm to be also sensitive to spindles in the deep sleep stage N3, in which the strong contribution of lower frequencies tends to reduce the relative sigma peak. This is especially relevant in the context of the known age-related differences in the low frequency power^83^, which may bias the algorithm towards more lenient detection in older adults. Lastly, spindles that were too close to each other (<500 ms) were merged, and spindles that were too short (< 500 ms) or too long (> 2 sec) were removed.

Slow oscillations (SOs) were also detected with the YASA software, based on algorithms from Massimini et. al. (2004)^84^ and Carrier et. al. (2011)^85^. We implemented a two-step procedure to apply participant-specific thresholds, adjusting for age-related differences in the slow oscillations power^43,83^. Data was bandpass filtered between 0.3 and 1.5 Hz. In the first step, no amplitude threshold was set, detecting all zero-crossings with pre-defined durations as potential SOs. Then, individual thresholds were calculated per person, so that the final SOs had to exceed 1.25 times of the negative peak and 1.25 times the peak-to-peak amplitude of all initial detections.

### Feature selection

Multistep procedure was applied for feature selection (Figure 2). First, 24 initial features were extracted (20 continuous and 4 categorical). These could be broadly classified into 4 themes: spindle-specific attributes (n=8), temporal context (n=6), post-spindle period variables (n=9), and topography (n=1). Then, variables were discarded based on multicollinearity (bivariate Pearson’s r > 0.7) and conceptual overlap. Lastly, some variables were discarded on a post hoc basis after initial analysis due to apparent biases (Supplementary materials 1). Altogether, 8 continuous and 3 categorical features were retained for the main clustering analysis.

**Figure 2.**
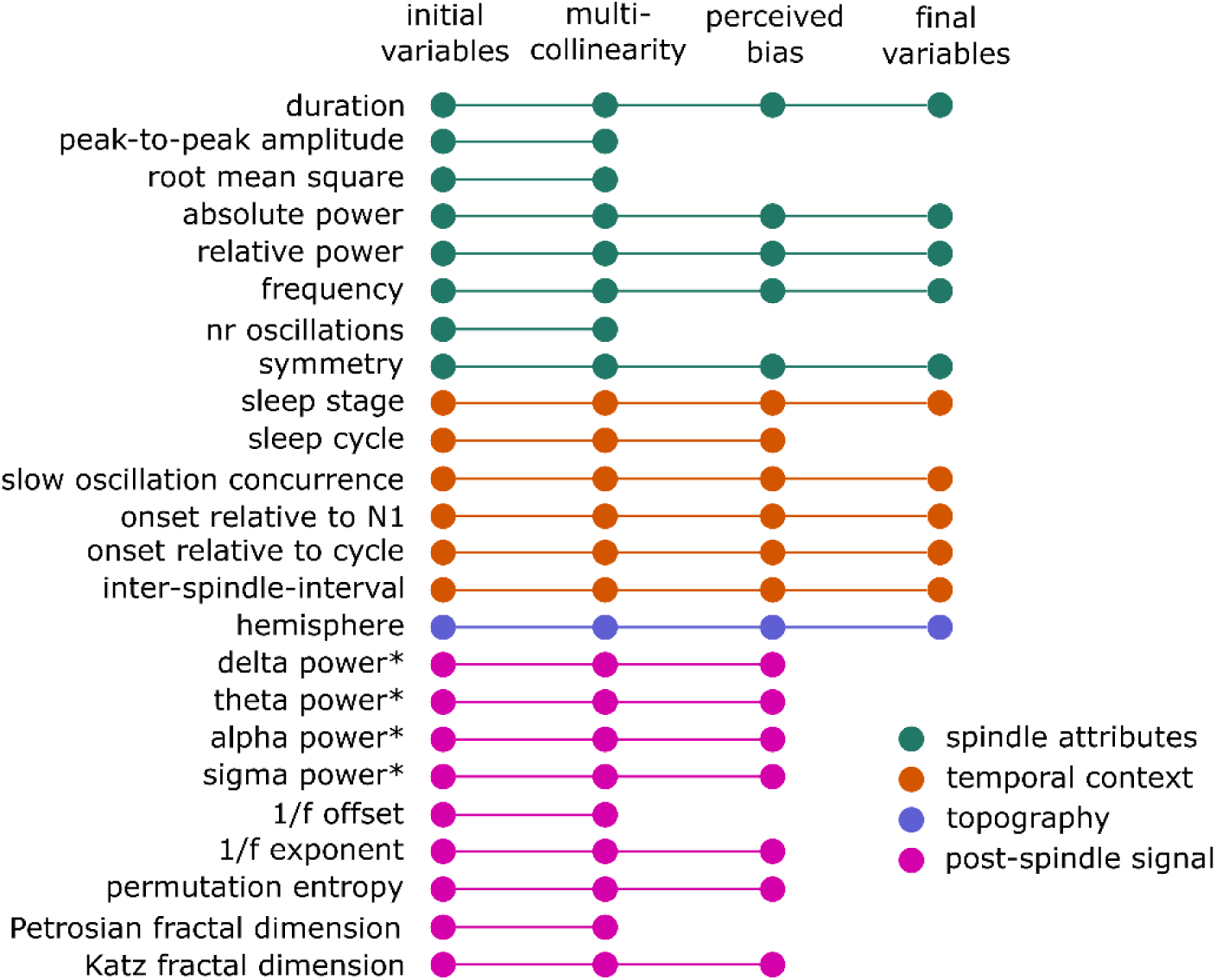
Selection of initial attributes and the multistep pruning of the attribute matrix. The colors of the nodes indicate attribute categories. Post-spindle signal attributes were extracted from the 2-second window after each spindle event. *marks relative power with respect to total broadband power spectrum.

The spindle-specific features were extracted from the YASA automatic output and included the duration, peak-to-peak amplitude, root mean square, median absolute power, median relative power, median frequency, number of oscillations, and symmetry. The absolute power and instantaneous frequency were derived from the Hilbert-transform of the sigma-filtered signal, and the relative power was derived from the short-term Fourier transform of the preprocessed signal, expressed as the percentage of power in the sigma band relative to the total power of the broadband signal. The symmetry indicates the location of the most prominent sigma peak, with 0 at the start and 1 at the end of the spindle, thus values around 0.5 indicating symmetric spindles.

A set of features were calculated reflecting the temporal context of each spindle. These included the stage in which the spindle occurred, sleep cycle, onset time relative to the sleep onset (first indication of N1 sleep), onset relative to the beginning of the cycle, inter-spindle interval, and co-occurrence with slow oscillations. Sleep cycle was determined by the adapted Feinberg’s criteria^86,87^ and only spindles within the first five cycles were retained. Spindles with an inter-spindle interval >3 min were discarded. The previously detected SOs were used to indicate their co-occurrence with the spindle. If the spindle start occurred within the time period between negative and positive peak of any of the detected SOs, the spindle was marked as occurring on the up-phase of the slow oscillation. Assuming symmetry of the slow oscillations, if a spindle occurred within the same time-window following the positive peak or preceding the negative peak, the spindle was marked as occurring on the down-phase. If neither of these conditions were met, the spindle was marked as not co-occurring with a SO.

The single topographical variable captured hemispheric distribution, as only electrodes C3 and C4 were available for all participants. Hemisphere was coded to indicate whether the spindle was detected in the left (C3), right (C4), or both hemispheres. The co-occurrence was determined if there was any overlap within the start and end time of the spindles in both electrodes, and only leading spindles were retained for the analysis. That is, if a spindle was detected simultaneously in both electrodes, only the spindle with an earlier start was included in the analysis.

The signal directly after the spindle may be informative regarding local cortical processing, thus the next set of features described the post-spindle period of two seconds. Welch’s method was used to calculate the power spectrum of the post-spindle period and relative power with respect to the total power was extracted for the canonic frequency bands in human sleep: delta (0.5-4 Hz), theta (4-8 Hz), alpha (8-12 Hz), and sigma (12-16 Hz). Next, the offset and the exponent of the frequency spectra were estimated using Python *fooof* package^88^ (v. 1.0.0). Lastly, the post-spindle signal complexity was captured by permutation entropy, Petrosian fractal dimension, and Katz fractal dimension, calculated with the Python *antropy* package^89^ (v. 0.1.6). These measures indicate the non-linear complexity of the signal, and can roughly be interpreted to reflect the predictability and regularity of the signal^90^, with higher complexity putatively indicating increased information processing^52,91^. Permutation entropy was chosen due it computational efficiency and Petrosian fractal dimension as it was ranking relatively high among the features for automatic sleep stage classification^78^. Katz fractal dimension was calculated, as it is considered more appropriate for detecting within participant state changes^92,93^

For all continuous variables, spindles +/-5 standard deviations were discarded to remove extreme outliers. Skewed variables (duration, amplitude, root mean square, relative power, number of oscillations, theta, alpha, sigma, onset to cycle, and inter-spindle interval) were log-transformed. Pearson correlation matrix was calculated across all continuous features and, of the pairs with r > 0.70, only one of the features were retained. Second pruning of the feature matrix was done after initial results were obtained (Supplementary Materials 1). These pruning steps were necessary to avoid artificially biasing the clustering algorithm towards features that conceptually capture the same phenomena, and to reduce the influence of sleep stage due to the inclusion of the post-spindle signal characteristics. However, as the post-spindle event attributes had a good a-priori explanation to be included and may reflect relevant processes for memory consolidation, we report the differences in these features between the age groups, as well as summarize the clustering results that included these variables in the Supplementary Materials 4.

### K-mean clustering

We applied a k-means clustering algorithm for mixed numeric and categorical data^94^, implemented in MATLAB v2023a. This algorithm builds upon the standard k-means approach by introducing a novel cost function and a distance measure suitable for mixed data types. For numeric features, the cluster centroid is the mean of all values, while for categorical features, it is represented as a proportional distribution of the values within the cluster.

Distances are calculated differently based on feature type: numeric attributes use weighted Euclidean distance, while categorical attributes rely on a co-occurrence-based distance function that reflects the distribution and relationships of category values across the dataset. The algorithm automatically determines the significance or weight of each feature, with more discriminative attributes contributing more to clustering. These adaptations allow the method to effectively partition datasets with mixed data types into meaningful clusters. For further details, refer to Ahmad & Dey (2007)^94^.

As all attributes need to be in the same units for unbiased distance calculation, all numeric values were normalized to range between zero and one. We chose to do this within each participant for greater generalizability of the results. That is, the normalized values for each participant are not affected by other participants’ values, so that that the cluster belonging of the spindles of any new participant could theoretically be calculated based on the cluster centroids of a representative sample. However, this normalization procedure reduces between-participant variability and therefore runs the risk of reducing associations with memory performance. Therefore, we repeated the clustering pipeline with the values normalized globally across all participants and summarize the results in the Supplementary Materials 4.

The clustering algorithm was run with several values for k, varying from k = 2-10. The final number of clusters was selected based on the visualization of the silhouette values and mean distances from cluster centroids. The maximum iterations per k was set to 300.

For visualization, we extracted 100 spindles with the lowest distance from the centroid per cluster and age group to represent the stereotypical spindle shape. This was done blind to participant identity, but the post-hoc examination indicated that 28-32 participants contributed to each cluster/age group. The spindle time-courses were bandpass filtered between 9-16 Hz, time-locked to the negative peak, zero-meaned, and averaged time-pointwise for visualization.

### Statistical testing

Statistical analysis was performed with R (v. 4.0.1).^95^

*Memory performance.* All six memory scores were entered into PCA to derive a composite measure. Independent t-tests were conducted to compare memory performance between old and young age groups. Baseline memory performance (AFC-1) was compared between sleep and wakefulness using linear mixed models with factors condition (sleep/awake), age group (old/young), and their interaction.

*Spindle attributes.* To fully characterize the spindle profiles before clustering and for comparability with previous literature, mean value per attribute and participant was computed. For categorical attributes, we calculated the percentage of spindles (1) belonging to N2 sleep, (2) co-occurring with SOs (regardless of up- or down-phase), and (3) bilateral topography. All features were compared between older and younger age-groups using independent t-tests and fdr-corrected p-values across all attributes.

As our clustering approach revealed two clusters, in which all or most spindles co-occurred with the SOs, we calculated the exact phase angle of the underlying SOs at the peak of the sigma amplitude for each spindle in the post-hoc basis. The details on statistical testing and results are listed in Supplementary Materials 5.

*Cluster profiles.* After clustering, we compared the distributions of the continuous features pairwise between all clusters using Kolmogorov-Smirnov tests and fdr-corrected p-values. However, due to the large number of spindles, all comparisons were significant and the statistical results are therefore not reported in detail. The clusters’ descriptions will be guided by the proportional distributions of categorical features and power density functions of the continuous features.

*Participants’ profiles in each cluster.* To characterize each participant’s expression of spindles in each cluster, we calculated two summary values per participant and cluster. Specifically, the density of each cluster characterizes the amount of spindles in that cluster per person, normalized to their sleep duration. Given that the clustering revealed spindle dissociation based on sleep stage, to prevent bias due to differences in the N2 and N3 durations, we normalized the densities with respect to the specific sleep stage that constituted each cluster. The second summary value represents the proportional distribution of spindles to clusters per person. That is, for each person, we calculated the percentage of spindles in each cluster from all this participant’s spindles.

The densities and proportional distributions were entered into separate mixed ANOVAs with factors cluster, age group, and sex, including their interactions. Significant effects were followed up by pairwise t-tests (independent for *age* and *sex* comparisons, and dependent for *cluster* comparisons) with Holm-Bonferroni corrections for multiple comparisons.

*Spindle attributes per participant and cluster.* To more specifically characterize the spindles in each cluster and age group, we extracted the averages (per participant and cluster) of the three most used spindle attributes in previous literature: spindle frequency, (absolute) power, and duration. These were compared using the same mixed ANOVAs and post-hoc tests as described above.

*Spindles’ associations with memory*. First, we ran sensitivity analysis to determine the minimal effect size we have power of 0.8 to detect, given our sample size. To test the associations between participants’ cluster profiles and age-related memory decline, we ran multiple regression models, predicting the combined memory performance (PC1) from the cluster score, age-group, sex, and their interactions. Separate models were run using density or proportional distribution as predictions, as well as power, duration, and frequency. In addition, to directly test the role of sleep in overnight memory consolidation, these models were repeated with the ‘sleep gain’ memory indicator as the outcome variable. All models were run separately for all clusters, so that with the selected k = 4, we considered a p-value 0.05/4 = 0.0125 as statistically significant.

## RESULTS

### Memory performance

Memory performance was quantified with six different measures (Table 1, Figure 3A). Age differences were evident in most measures, except for the experimental sleep gain. Whereas the experimental memory scores were higher in younger participants, unexpectedly, they reported more subjective problems on the everyday memory questionnaire than the older adults.

**Figure 3.**
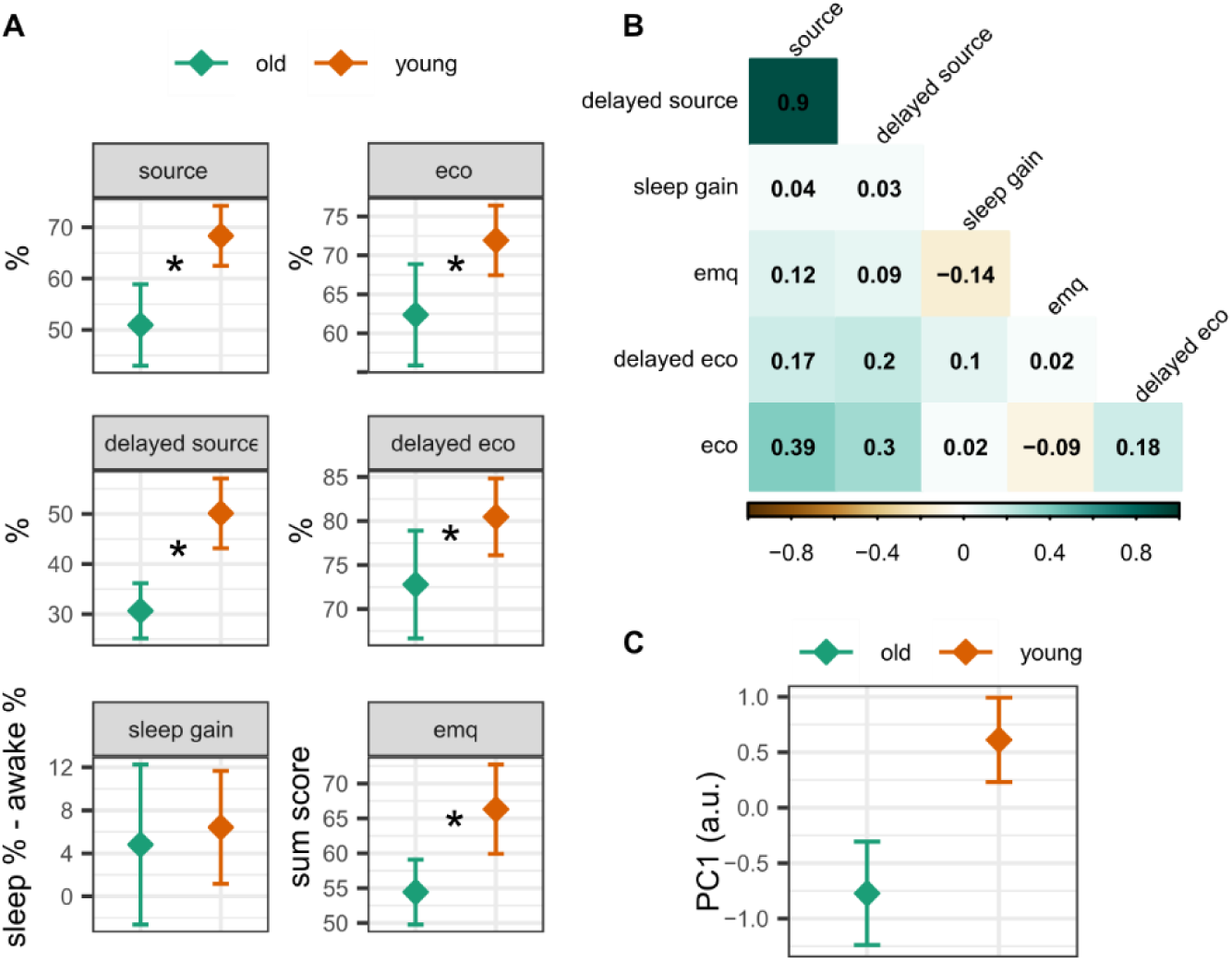
(A) Memory performance per age group (young n = 43, old n = 34), separately for each measurement. The error bars represent 95% confidence intervals around the mean. (B) Correlation matrix between different memory scores. (C) Age differences in the first principal component scores. *indicates significant differences between the young (n=43) and old (n=34) age group.

**Table 1.** Memory performance and statistical comparisons between young (n = 43) and old (n = 34 age groups. The values for young and old are mean values, with standard deviations in the brackets. *indicates significant p-value at the alpha level of 0.05.

| Memory score | Explanation | old<br>mean (sd) | Young<br>mean<br>(sd) | t (df) | Cohen's d | p-value |
| --- | --- | --- | --- | --- | --- | --- |
| source recall | Experimental retrieval performance (%) after 12 hours | 51 (24) | 68 (20) | -3.45 (64) | -0.81 | 0.001* |
| delayed source recall | Experimental retrieval performance (%) after ~6 days | 31 (16) | 50 (23) | -4.30 (74) | -0.95 | <0.001* |
| sleep gain | Difference between source recall (5) between sleep and awake conditions | 5 (22) | 6 (18) | -0.35 (62) | -0.080 | 0.730 |
| ecological recall | Ecological memory test performance (%) at the end of the session | 62 (19) | 72 (15) | -2.37 (61) | -0.56 | 0.021* |
| dealyed ecological recall | Ecological memory test performance (%) four weeks later | 73 (18) | 80 (15) | -2.00 (62) | -0.470 | 0.049* |
| emq | Everyday memory questionnaire | 54 (14) | 66 (21) | -2.94 (72) | -0.640 | 0.004* |

The sleep gain effect was calculated as the difference in source memory scores between sleep and wakefulness conditions. Both younger and older adults recalled 5-6% more item-category associations after a 12-hour sleep period compared to wakefulness. Separate results for the sleep and awake conditions across different retrieval tests in the source memory experiment can be found in Supplementary Materials 2 (Table S1). It is noteworthy that the values between sleep and wakefulness correlated highly, with correlations ranging from r = 0.72 to 0.76. Analyzing the conditions separately using a linear mixed ANOVA revealed a main effect of condition (sleep > awake, F = −2.71, p = 0.024) and age group (young > old, F = −8.82, p < 0.001), indicating a similar sleep gain over wakefulness for both age groups (non-significant interaction; F = 0.50, p = 0.676).

The subtractive sleep gain effect was not adjusted for baseline performance, as the exact category-level retrieval test (differentiation between faces and places without visual cues) was not administered immediately after encoding. Instead, participants completed a forced choice test, selecting the specific face or place for each item (AFC-1, Figure 1). In this context, a mixed linear model revealed a significant effect of age group (F = −7.98, p < 0.001) but no condition (F = −1.52, p = 0.12) or interaction effects (F = 0.24, p = 0.80), indicating no significant baseline learning differences between sleep and awake conditions (see Table S1 for separate values for sleep and awake conditions). Nevertheless, we recalculated the sleep gain measure using forced choice performance relative to baseline. This analysis confirmed greater retention after sleep compared with wakefulness (young: mean = 5%, sd = 11%; old: mean = 9%, sd = 14). This adjusted measure strongly correlated with the original sleep gain measure used in the PCA (r = 0.69).

The correlation structure of the six main memory scores are depicted on Figure 3B. The experimental 12-hours source memory recall correlated strongly with the delayed source memory (r = 0.90), and these two correlated moderately with the immediate ecological memory test (r = 0.30 – 0.39). Otherwise, correlations between the scores were negligible.

The first PC (37% variance explained, Figure 3C) combined the age differences across all six memory scores (t(68) = −4.51, Cohen’s d = −1.04, p < 0.001), whereas no age differences were found between PCs 2-6 (all Cohen’s d-s <= |0.37|, all p-values >= 0.119). The factor loadings indicated strongest contributions to PC1 from the 12-hours and 6-days experimental source recall, and additional modest contributions from the immediate and four weeks ecological recall. The contribution of the sleep gain effect and everyday memory questionnaire to PC1 were negligible. Given that the age differences in memory performance were captured by PC1 alone, this will be used for further testing associations between age-related memory performance and sleep spindles. However, since the sleep gain measure may more specifically reflect sleep-facilitated memory consolidation, this will be tested separately, despite no age differences being present in this measure.

### Sleep architecture and spindles

Participants slept six hours on average (sd = 0.67) on the experimental night, with older participants having shorter total sleep durations, shorter proportion of N3 sleep, and higher proportions of N1 and N2 sleep, in accordance with previous observations of reduced deep sleep in older adults (Supplementary materials 3, Table S2). Subjective sleep problems, measured by Pittsburg Sleep Quality Index global score, did not differ between older and younger participants (mean = 5, sd = 2, for both young and old, t(62) = −0.23, d = −0.05, p = 0.819).

Across all participants, 99’188 spindles were used in the analyses. Of these, 57’614 belonged to the younger and 41’574 to older participants. On average, each participant contributed with 1’288 (sd = 473) spindles, and average spindle densities were 5.44 per minute of NREM stage N2 and N3 sleep. There were no differences between old and young age groups in average spindle counts and densities. Note that we have slightly higher spindle densities as commonly reported in previous literature, likely due to differences in the automatic spindle detection parameters. First, we included a broad frequency range of 9-16 Hz, incorporating both fast and slow spindles. Second, we discarded a commonly used relative power thresholding to be sensitive towards stage N3 spindles, given the lower sigma peak relative to low frequencies during deep sleep.

### Spindle attributes per age group

The summary values for all features per age group are presented in Table 3, together with details on statistical tests. The differences between old and young groups in spindle attributes can be summarized as follows: older group had lower absolute power and dominant frequencies. The duration, relative power, and symmetry were similar across age groups. Older adults had lower proportion of bilateral spindles, compared with younger age group.

**Table 3.**
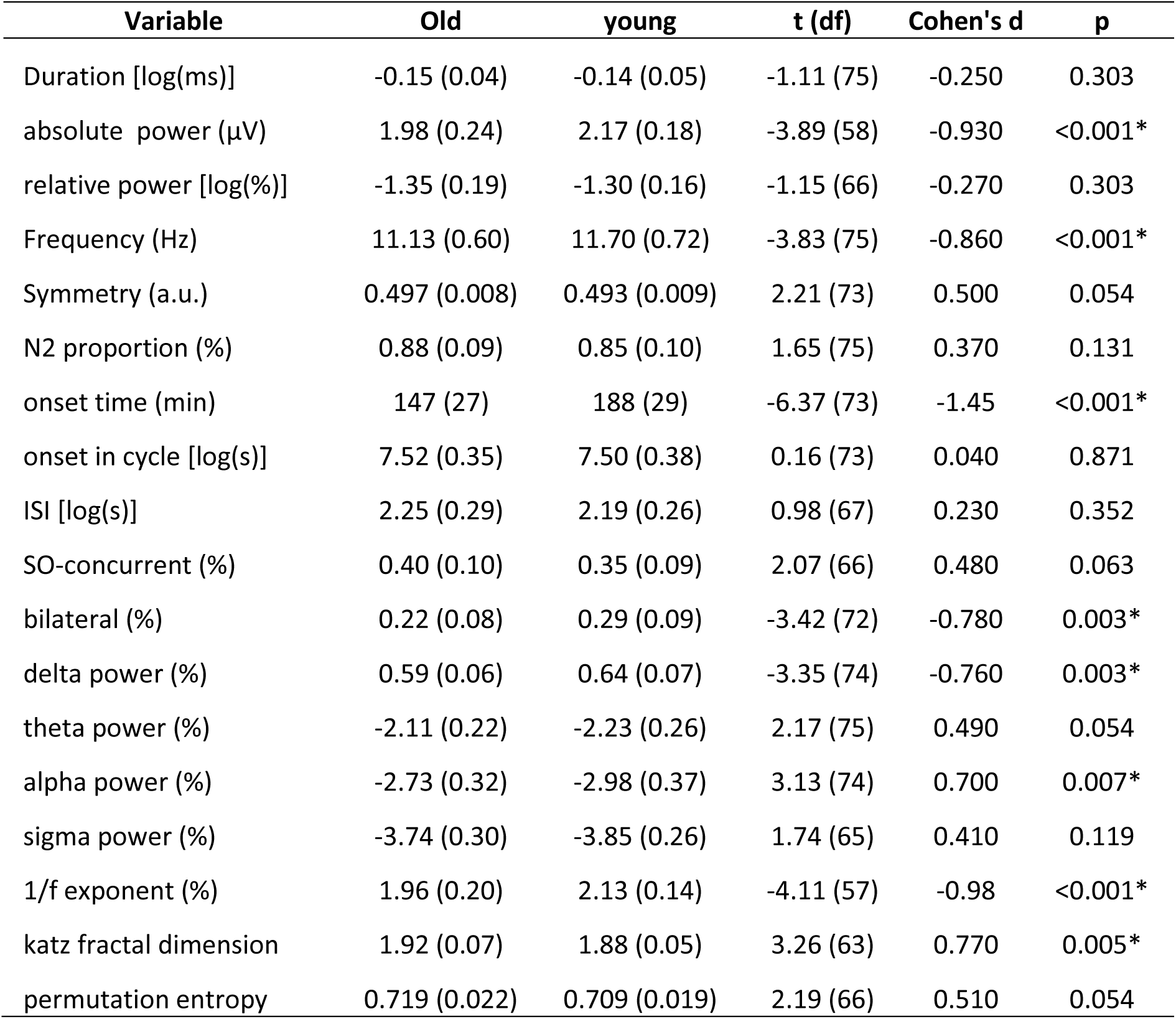
Average values (and standard deviations) for spindle attributes per age group, together with details about statistical testing. Young n = 43, old n = 34. *marks a fdr-corrected significant difference at an alpha value of 0.05. ISI = inter-spindle-interval, SO = slow oscillation

Regarding the temporal context, older participants’ spindles tended to occur earlier in the night, likely due to younger participants sleeping longer and having more opportunities for spindle activity in the morning hours. There were no statistical differences regarding the proportional distribution of spindles with respect to sleep stage or their co-occurrence with SOs, nor between the inter-spindle interval or onset time relative to the start of the cycle.

Regarding the post-spindle signal characteristics, the older age group had lower delta power, lower alpha power, smaller 1/f exponent, and higher Katz fractal dimension. Altogether, these findings reflect higher proportion of slow frequency activity in younger people, together with higher post-spindle signal complexity.

### Cluster selection and feature significances

We explored different cluster solutions, varying the number of clusters k = 2-9. The solution with k = 4 had the highest average silhouette value and a there was a negligible decrease in average distances after k > 4 (Figure 4A). The silhouette scores for k = 4 are visualized in Figure 4B. In an ideal case, the silhouette values in each cluster would have values close to one and rectangular cluster shapes instead of smooth curves, and no negative values. This was not the case for any k, indicating a continuity of spindle features, rather than distinct separability.

**Figure 4.**
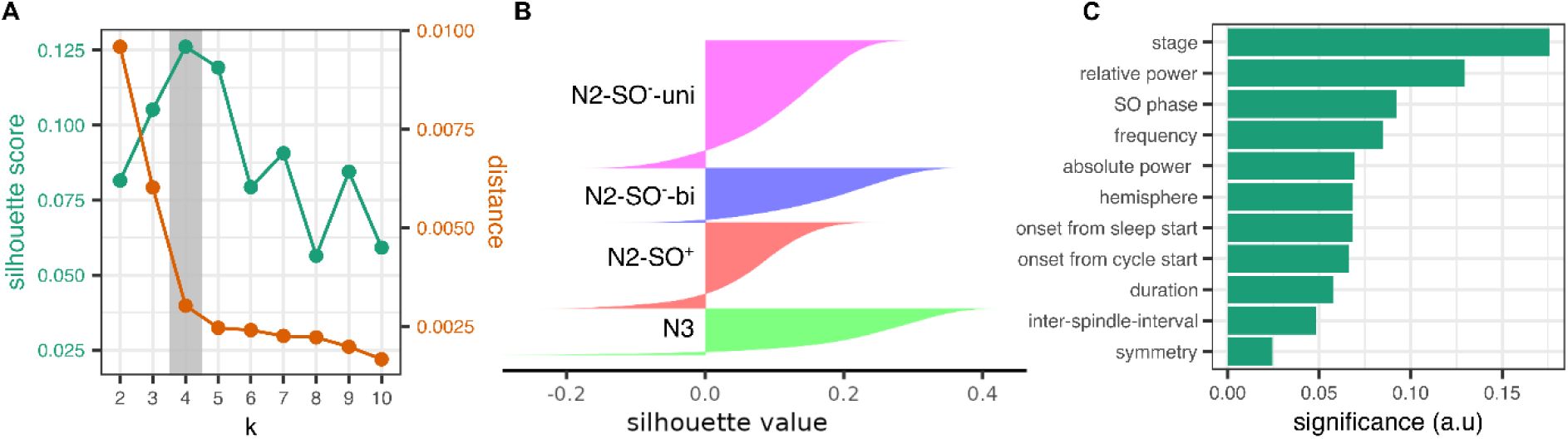
(A) Mean silhouette values (green) and mean distance from the cluster centroid (orange) across all spindles as a function of k. The shaded are represent values at k=4, which was selected for further analysis.(B) Silhouette values for the spindles in each cluster. (C) Significance of each attribute to the separability of the clusters, as determined by the co-occurrence of values in the data, common across all k’s.

The feature significances are depicted at Figure 4C and indicate that sleep stage had the highest contribution to clusters’ separability, followed by relative power, slow oscillation concurrence, and spindle frequency.

### Clusters’ descriptions

The visualization of the spindle time courses, averaged over 100 spindles with the lowest distance from their respective centroids, are depicted on Figure 5A. Intuitively, these represent the stereotypical spindle shapes for each cluster and age group. With k = 4, the combination of categorical features fully distinguished each cluster, which guided cluster naming henceforth. Stage N3 spindles formed a distinct cluster (15% of all spindles), as did stage N2 spindles that co-occurred with SOs (27%; henceforth named N2-SO^+^). The remaining N2 spindles were subdivided into those with unilateral (41%) and bilateral (17%) topography (named N2-SO^-^-uni and N2-SO^-^-bi, respectively). With the alternative pipelines using the group-normalized data or including post-spindle signal features, the results were qualitatively similar (Supplementary Materials 4). In the following, each cluster is described in detail, based on the proportional distribution of categorical features (Figure 5B) and power density functions (Figure 5C). As symmetry and absolute power had smaller discriminative differences, these features will not be discussed in detail. We also summarize the results from the SO-coupling for the N3 and N2-SO^+^ cluster (see Supplementary Materials 5 for the specific statistical tests for the SO-coupling).

**Figure 5.**
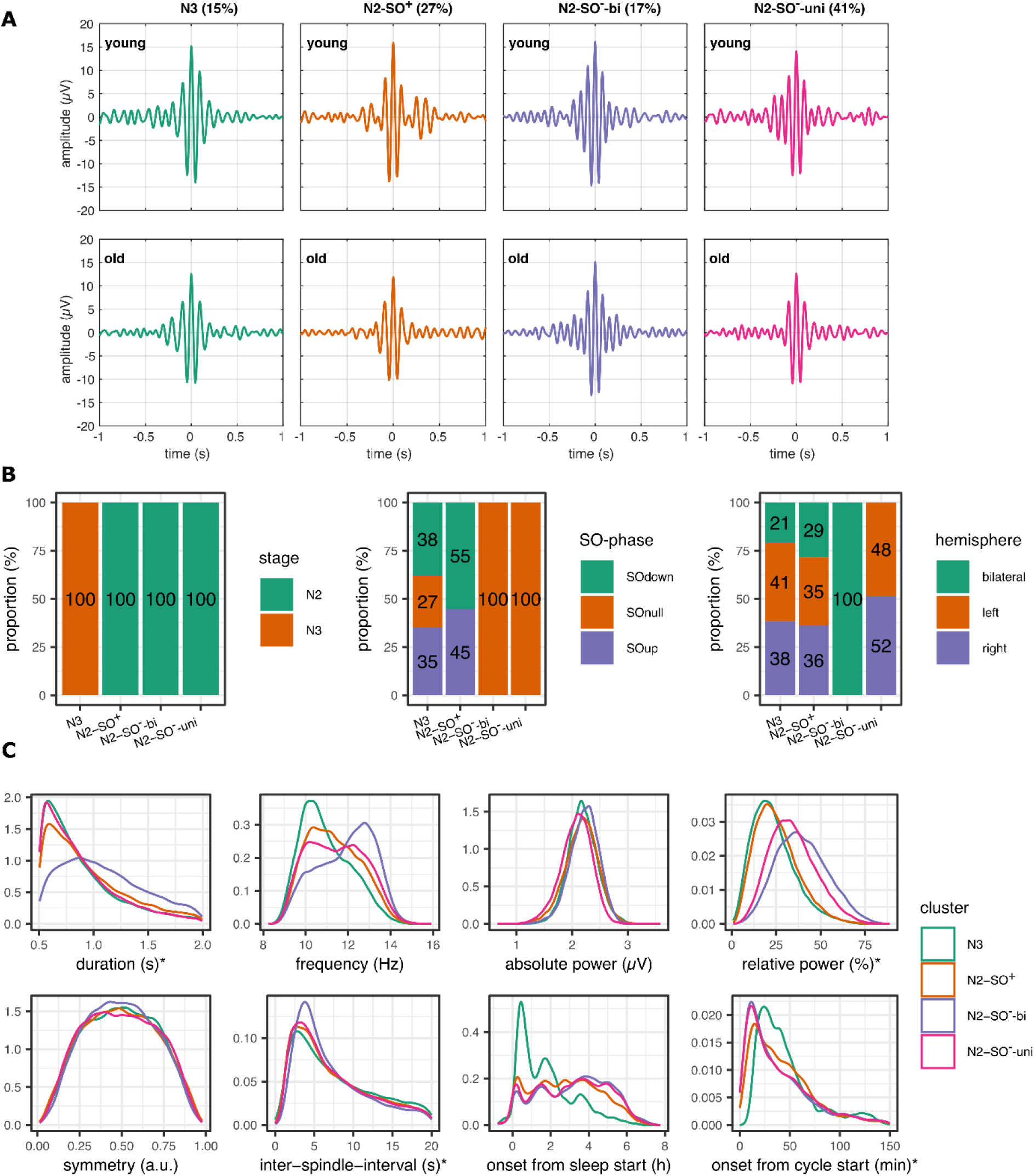
(A) Stereotypical spindle time-courses for each cluster and age group, averaged over 100 spindles with lowest centroid distance in each group. The percentage indicates the proportion of the spindles in the cluster of all spindles, regardless of age group. (B) Proportional distribution of categorical variables within each cluster. (C) Power density functions of continuous variables within each cluster. In total, number of spindles that are used for the calculation of the density functions was 57 614 for the young, and 41 574 for the old age group. Please refer to table 3 for the number of spindles in each cluster. *marks variables, which were log-normalized in clustering due to skew, but are depicted here in their original units for better interpretation.

*The N3 cluster* formed the smallest cluster (15% of all spindles) and was dominated by spindles co-occurring with the SOs (73%). Both uni- and bilateral spindles were collapsed in this cluster. The N3 spindles had a frequency distribution towards the slower range, peaking at around 10.5 Hz, shorter durations, and lower relative power. They were distributed earlier during the night, reflecting the increased proportion of slow-wave sleep in the first half of the night. In contrast, they appeared later in each cycle, reflecting the natural order of sleep stage N2 preceding N3 in each cycle. In young adults, they were distributed uniformly on the SO up-phase with a lower-than-chance occurrence on the down phase, whereas in the older adults, they occurred preferentially around the negative peaks of the SOs (Figure S1).

*The* N2-SO^+^ *cluster* (27%) represents those spindles that occurred in stage N2 and co-occurred with the SOs. Interestingly, the data-driven clustering algorithm only differentiated whether the spindles coincided with the SO or not, with no consideration of whether the spindle occurred in the up or down phase. However, the post-hoc SO-sigma phase-amplitude coupling indicated preferred occurrence just before or at the positive peak in young adults, while the distribution was shifted towards the SO negative peaks in older adults (Figure S1). The spindles in these cluster occurred both uni- and bilaterally, were distributed uniformly across the night, but occurred slightly later in each cycle than other N2 spindles. Their frequency distribution was biased towards the lower range, with a broad peak around 10-11 Hz.

*The* N2-SO^-^-uni was the largest cluster (41%), consisting of stage N2 unilateral spindles that did not co-occur with SOs. These were distributed uniformly through the night and had bimodal frequency distribution peaking at around 10 and 12 Hz. They had relatively higher power, compared with stage N3 and to the N2 spindles, which co-occurred with the SOs.

*The* N2-SO^-^-bi *cluster* (17%) consisted of stage N2 bilateral spindles that did not co-occur with SOs. This cluster stood out, as it had the highest relative power, longest inter-spindle-intervals, longest durations, and highest central frequencies, peaking at around 13 Hz.

### Expression of cluster-specific spindle activity

To compare the expression of resulting sleep spindle clusters between age groups, we calculated spindle density and proportion per participant and cluster (Figure 6). The correlations between densities in different clusters ranged between r=0.11 to r=0.65, with the highest correlation between N2-SO^-^-uni and N2-SO^-^-bi. The correlations between percentages in different clusters were smaller, ranging between r=-0.06 to r=-0.57. Therefore, the densities and percentages between clusters were not redundant. For example, a person with high density in one cluster did not necessarily have high density in a different cluster.

**Figure 6.**
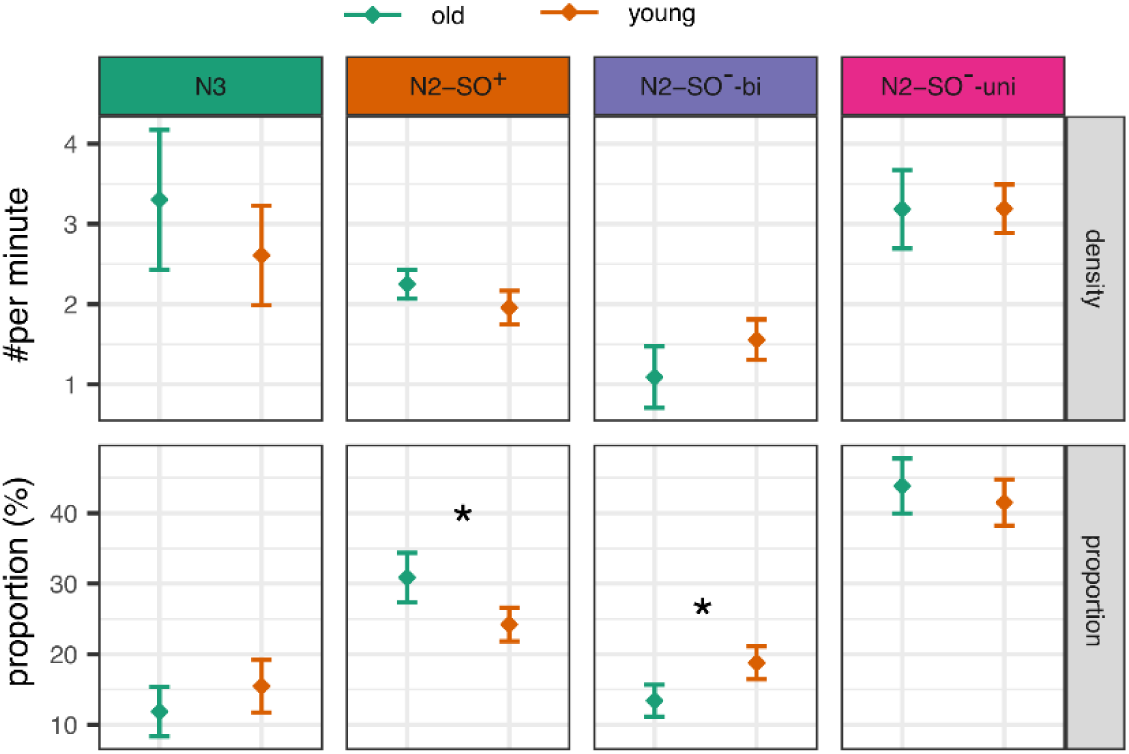
Expression of spindle clusters, contrasting younger (n=43) and older (n=34) age groups. The error bars represent 95% confidence intervals, corrected for within-participants measurements between clusters. *marks variables with significant differences between age groups, given a statistically significant interaction between age group and cluster.

*Density.* Spindle densities indicate the number of spindles per minute of sleep stage they occurred in. That is, given that the clustering results revealed perfect dissociation between stage N2 and N3 spindles, they were normalized to those durations, respectively, not to bias density values towards differences in sleep stage durations. Stage N2-unilateral spindles had highest densities (mean = 3.19 per minute, sd = 0.94), followed by N3 spindles (2.92 per minute, sd = 2.69), then N2-SO spindles (2.09 per minute, sd = 0.86), and finally N2-bilateral spindles (1.35 per minute, sd = 0.76; main effect of *cluster*: (F(1.21, 88.46) = 27.99, η_p_^2^ = 0.28, p < 0.001). All post-hoc pairwise comparisons between cluster densities were statistically significant (all Cohen d’s >= 0.32 all p’s <= 0.004), except between the two densest clusters (N3 vs. N2-SO^-^-uni: Cohen’s d = 0.14, p = 0.412). There were no significant age or sex differences, nor interaction effects, indicating similar densities for younger and older age groups (all η_p_^2^’s <= 0.04, all p’s <= 0.101).

*Proportional distribution.* The proportional distribution indicates, which cluster dominated for each person throughout the night. Note that these are not independent from each other – as the proportion of one cluster increases, it must come at the expense of another. Both age groups followed the distribution of the overall spindle distribution, with highest proportion of N2-SO^-^-uni spindles (mean = 43%, sd = 9), then N2-SO^+^ spindles (mean = 27%, sd = 8), then N2-SO^-^-bi spindles (16%, sd = 7), and finally N3 spindles (mean = 14%, sd = 10; main effect of cluster: F(2.56, 186.98) = 15.03.99, η_p_^2^ = 0.67, p = < 0.001). All pairwise comparisons were significant (all d’s >= |1.43|, all p’s < 0.001), except the N3 and ‘N2-SO^-^-bi percentages (d = −0.30, p = 0.080).

There was no main effect of age on the distributions (F(1, 73) < 0.01, η_p_^2^ < 0.01, p ∼ 1). However, an interaction between age and cluster indicated cluster-specific age differences (F(2.56, 186.98) = 7.44, η_p_^2^ = 0.09, p < 0.001), which were further modulated by sex (three-way interaction F(2.56, 186.98) = 3.17, η_p_^2^ = 0.042, p = 0.003). Specifically, older adults had a higher proportion of N2-SO^+^ spindles, compared with the younger age group (t(75) = 3.77, d = 0.87, p = 0.001). After correction for multiple comparisons, this comparison only remained significant in men (men: t(32) = 3.44, d = 1.20, p = 0.012; women: t(41) = 2.04, d = 0.62, p = 0.190). These differences were reversed in ‘N2-SO^-^-bi spindles, with older adults showing lower proportion (t(75) = −3.80, d = −0.87, p = 0.001). Again, when compared separately in men and women, the age group comparison only remained significant for men after multiple comparison correction (men: t(32) = −3.61, d = −1.26, p = 0.008; women(t(41) = −0.72, d = − 0.72, p = 0.106). There were no age differences in N3 (t(75) = −1.62, d = −0.37, p = 0.219) and N2-uni-SO spindles (t(75) = 1.08, d = 0.25, p = 0.282).

In sum, there was a proportional drift in spindle expression in older adults, showing lower proportion of N2 bilateral spindles that were separate from the SOs, and a comparative increase in the proportion of N2 spindles that co-occurred with the SOs. These differences were more pronounced in men than in women.

### Participant- and cluster-specific spindle attributes

The spindle shapes in Figure 5A hint differences in spindle attributes between clusters and age groups. We therefore calculated averages per participant and cluster for the three most used spindle attributes in the literature: frequency, (absolute) power, and duration (Figure 7). However, there were very high correlations between the clusters with respect to frequency (ranging between r = 0.82 to r = 0.97) and power (between r = 0.92 to r = 0.99). That is, a person who had a lower frequency and/or power in any one cluster, also had lower values in all other clusters. Correlations between spindle durations between clusters were more modest, ranging between r = 0.40 to r = 0.63.

**Figure 7.**
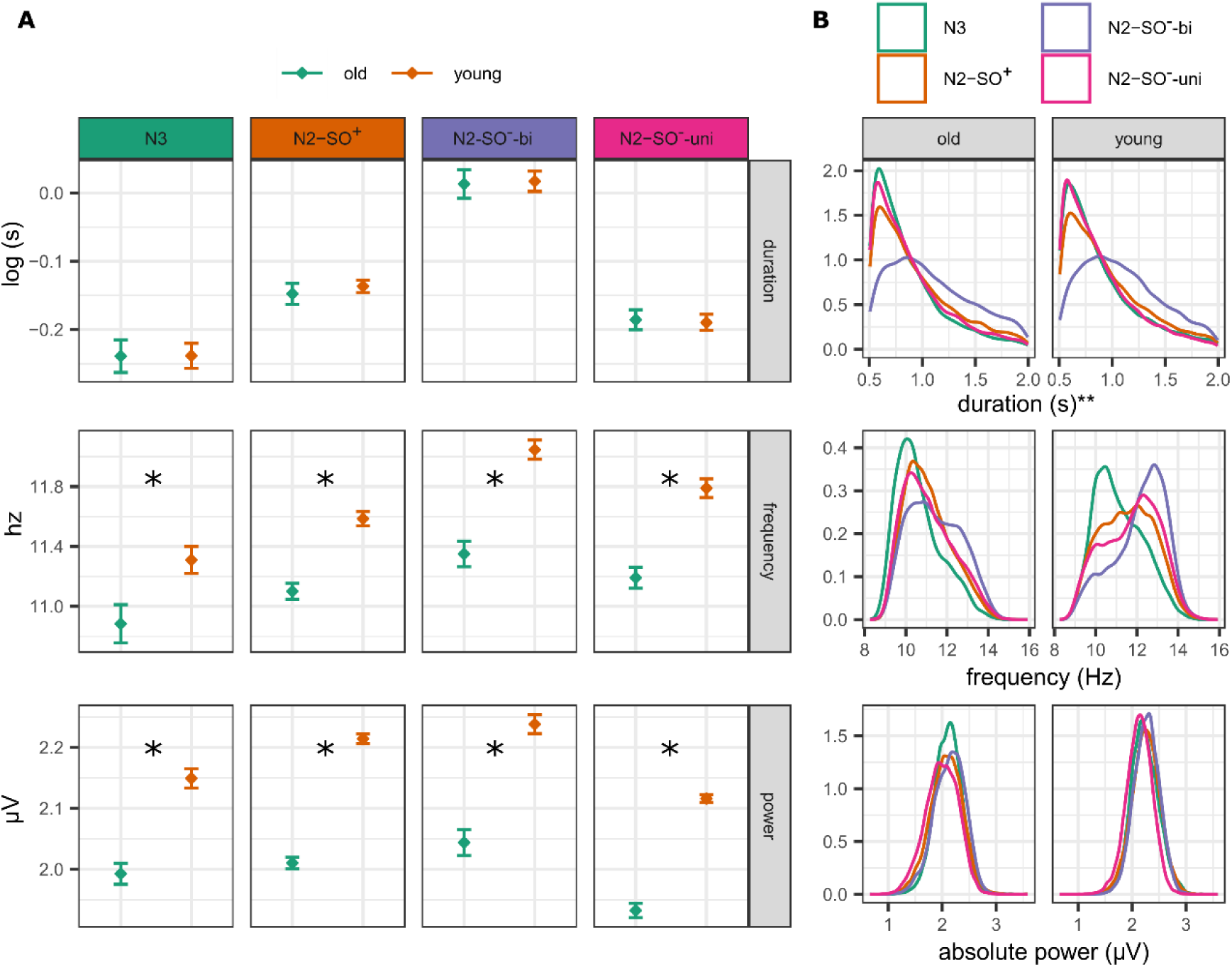
(A) Average duration, frequency, and power across participants, separately for each cluster and age group (young n = 43, old n = 34). The error bars represent 95% confidence interval, corrected for within-between measurement across cluster. (B) Power density profiles for each cluster and age group for duration, frequency, and power. In total, number of spindles that are used for the calculation of the density functions was 57 614 for the young, and 41 574 for the old age group. Please refer to table 3 for the number of spindles in each cluster. * marks significant difference between age groups; ** marks that the variable was log-normalized for clustering and statistical testing, but is here represented in original units.

*Frequency*. There were considerable frequency differences between the clusters (F(1.73, 126.23) = 83.49, η^2^_*p*_ = 0.53, p < 0.001), as well as age differences (F(1, 73) = 13.2, η^2^_*p*_ = 0.15, p < 0.001). Whereas there was also a significant interaction of age and cluster interaction (F(1.73, 126.23) = 5.75, η_p_^2^ = 0.07, p = 0.006), all pairwise comparisons still indicated higher frequencies in younger than older in each cluster (all d’s >= |0.93|, all p’s <= 0.016). Similarly, differences in frequencies between clusters in both age groups were all significant (young: all d’s >= 5.64, all p’s <= 0.001; old: all d’s >= |2.15|, all p’s <= 0.039). Specifically, older adults had the biggest frequency difference in the N2-SO^-^-bi cluster, compared with the young age group. This shift in frequencies is starkly illustrated in the probability density functions over all spindles, divided by age group (Figure 7B). Whereas the N2-SO^-^-bi cluster has a sharp high-frequency peak in the young age group, this peak is shifted towards the lower frequencies in older adults.

*Power.* There were considerable cluster (F(1.79, 230.9) = 108.93, η_p_^2^ = 0.60, p < 0.001), age (F(1, 73) = 21.80, η^2^_*p*_ = 0.23, p < 0.001), and sex (F(1, 73) = 16.69, η^2^_*p*_ = 0.18, p < 0.001; women > men) differences in absolute spindle power, superseded by significant cluster and age interaction (F(1.79, 230.9) = 4.47, η_p_^2^ = 0.06, p = 0.016). Despite this significant interaction, the power was higher in the younger age group than older in all clusters (all d’s >= |0.74|, all p’s <= 0.002). Generally, the power was the highest in the N2-SO^-^-bi cluster, followed by the N2-SO^+^, N3, and finally the N2-SO^-^-uni cluster. All pairwise comparisons were significant in both age groups (young: all d’s >= |0.16|, all p’s < 0.001; old: all d’s >= |0.13|, all p’s <= 0.009), except for between N2-SO^+^ and N3 cluster in the older adults (d = 0.07, p = 0.008). In sum, power was larger in young than old age group in all clusters. However, power differences between clusters were generally stronger for the young than for the older age group.

*Duration*. The spindle durations were longest in all the stage N2 clusters (‘N2-SO^-^-bi > N2-SO^+^ > N2-SO^-^ -uni > N3; main effect: F(2.27, 165.72) = 355.88, η_p_^2^ = 0.83, p < 0.001; pairwise comparisons: all d’s >= |0.73|, all p’s < 0.001). There were no age (F(1, 73) = 0.15, η_p_^2^ = 0.002, p = 0.705) or sex differences (F(1, 73) = 3.35, η_p_^2^ = 0.04, p = 0.071).

### Clusters’ associations with age-related memory performance

The memory associations were tested with linear models, predicting memory performance (PC1) from cluster value, age group and their interactions, controlling for sex. PC1 was selected as a combined memory outcome measure, as this captured the age-related variance across all memory indices. In our sample, we had 0.8 power to detect R^2^ of 0.14. The sensitivity analysis for individual predictors (one-sample t-test) indicated that we had 0.8 power to detect an effect size of d = 0.32 in an uncorrected alpha level of 0.05, or d = 0.39 in an alpha level of 0.0125, which is the corrected alpha for separate models over four clusters.

The models predicting PC1 from spindle density or proportional distribution all had R² ≥ 0.30. Since these models were adjusted for age group and sex, the high explained variances are attributable to these covariates. Regarding the effects of interest, the proportion of the N2-SO^-^-bi cluster significantly interacted with the age group (b = 0.06, standardized beta = 0.71, p = 0.011). Separate linear regression analyses for young and older groups, controlling for sex, indicated a significant negative association in the younger group, with a higher proportion of N2-SO^-^-bi spindles associated with lower memory performance (Figure 8A). A similar trend was observed in spindle densities for this cluster (interaction with age: b = 0.46, standardized beta = 0.46, p = 0.026), although this did not reach significance at the corrected alpha level. The proportions and densities of the other clusters did not predict age-related memory decline, nor did any cluster-specific spindle attributes (frequency, power, duration; all standardized beta coefficients ≤ 0.19, all p-values ≥ 0.102).

**Figure 8.**
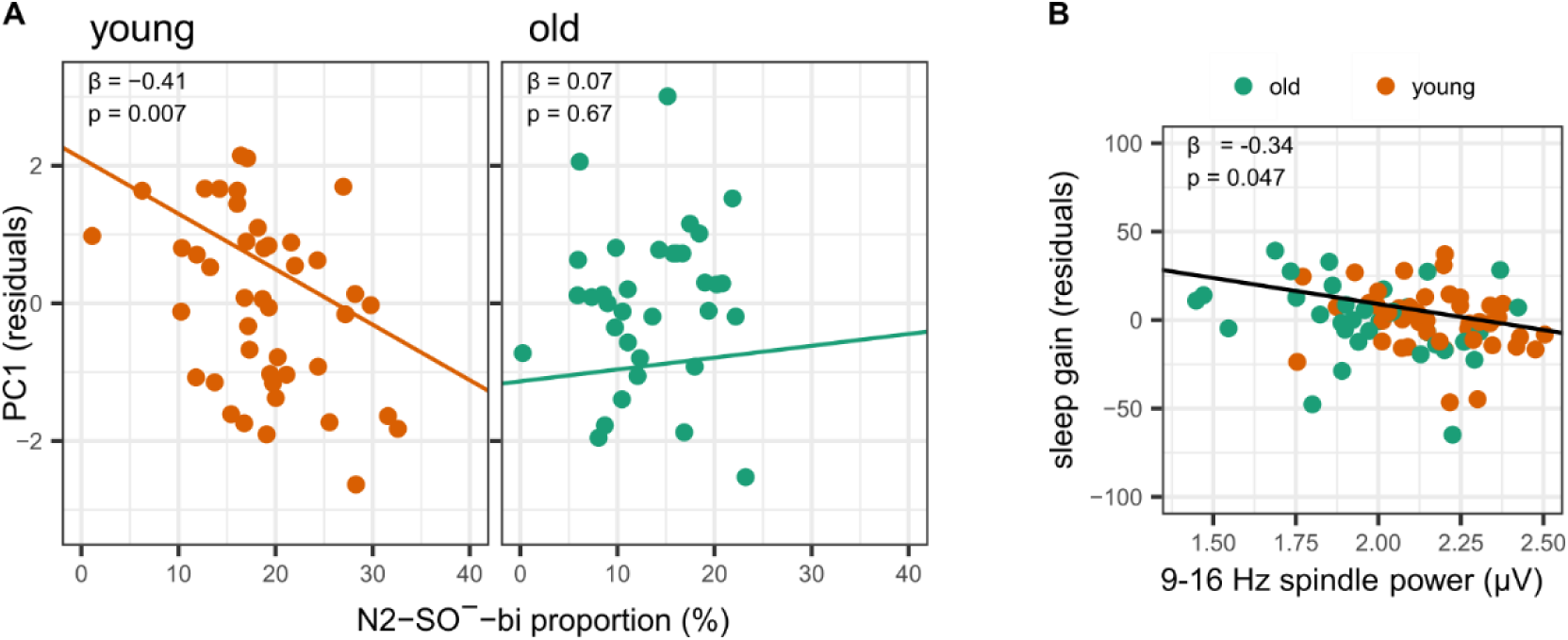
(A) Associations between age-related composite memory performance (PC1) and the proportion of cluster N2-SO^-^-bi spindles, indicating the interaction with age group. (B) Associations between sleep gain and overall spindle power (9-16 Hz), regardless of age group. The memory scores are residualized for sex (and age group in B), but the regression lines and text annotations represent the intercept, slope, beta, and p-values from the original models.

For completeness, we ran the same models using the spindle features (duration, absolute power, relative power, frequency, symmetry) calculated globally over the 9-16 Hz range, as well as total spindle densities, but found no significant associations with age-related memory performance even at an uncorrected alpha level.

### Clusters’ associations with the sleep gain effect

For the age-related memory associations, we used a combination measure over several different memory estimates. This included a custom ecological memory test. Whereas it was designed specifically to capture memory functioning in everyday activities, it has not been commonly used, is not standardized, and may add noise due to the narrative-based transcription procedure. This is partially evident in the higher scores in the delayed test, compared with the immediate, which likely derives from practice effects and higher number or items that were difficult to transcribe (deemed ‘irrelevant’ and discarded from the analysis) from the delayed narration four weeks later. In addition, we included the Everyday Memory Questionnaire, which unexpectedly indicated more subjective memory problems in the younger age group than in the older. This may be due to reporting bias or busier schedules of the younger participants with more chances for errors, indicating that this may not be the best instrument for assessing differences between groups with different lifestyles. We therefore repeated the analysis, using only the sleep gain effect, captured by the difference in memory performance after sleep, compared with the equivalent period of wakefulness. Note that this measure did not have age differences to start with. We found no significant memory effects for cluster densities and proportional distributions using this variable. Notably, the R^2^ of all models dropped to <0.07, likely since age group no longer was a significant predictor. However, we found associations for the absolute power in all cluster models (standardized beta values −0.32 to −0.40, uncorrected p-values <= 0.017; Figure S2). Although this relationship was significant at the corrected alpha level only in two clusters (N2-SO^+^ and N2-SO^-^-bi), the power was highly correlated between clusters, and therefore likely reflecting spindle power and memory associations regardless of the cluster belonging. We therefore additionally predicted the sleep consolidation effect from the general spindle power (that is, averages across all participant’s spindles from 9-16 Hz) and found a significant association (Figure 8B; b = −29.38, standardized beta = −0.34, p = 0.047), in which participants with higher spindle power had reduced sleep gain, when adjusted for age group differences. This relationship remained qualitatively similar if sleep gain was calculated on the forced choice test performance relative to the baseline performance (for cluster-specific models, all beta’s between −0.27 to −0.33, p-values between 0.029 to 0.064; for all spindles between 9-16 Hz, beta = −0.39, p = 0.029). Other spindle-related variables (duration, relative power, frequency, symmetry) were not associated with the sleep gain effect.

## Discussion

By leveraging spindle heterogeneity throughout the night, we identified four spatiotemporal spindle clusters with varying attributes and differential expression in aging. Notably, these spindles were clustered based on sleep stage, slow oscillation (SO) coupling, and hemispheric involvement (unilateral vs. bilateral). Among the older adults, there was a reduced proportion of N2 bilateral spindles, which was compensated by an increase in N2 spindles coupled with SOs. No density or proportional differences were observed in stage N3 and N2 unilateral spindles. Despite demonstrating non-random spindle heterogeneity and cluster-specific differences in spindle expression with age, the continuous variation of spindle attributes complicates the interpretation of the clusters as functionally distinct spindle types. Instead, the spindle heterogeneity better aligned with global signal differences between different NREM sleep stages.

### Parsing spindle heterogeneity

The categorical attributes – sleep stage, SO concurrence, and hemisphere – defined the final clustering solution hierarchically. First, all stage N3 spindles formed a distinct group, irrespective of SO concurrence or hemispheric involvement. Next, all stage N2 spindles that coincided with SOs formed a separate cluster, regardless of hemisphere. Finally, the remaining stage N2 spindles were divided into unilateral and bilateral clusters. The latter is reminiscent of a recent clustering attempt based on whole-head topography, which identified one global and several locally expressed clusters^41^. While our approach cannot precisely determine the topographical range of these emerged clusters, the bilateral spindles demonstrate the capacity to propagate across hemispheres, at a minimum.

In addition to categorical features, spindles within each cluster significantly differed in attributes such as frequency, duration, and relative power. The bilateral stage N2 spindles were notable for having the highest frequencies and longest durations, potentially corresponding to the ’fast spindles’ distinction in the literature. However, consistent with previous clustering work^41^, we observed wide frequency distributions across all clusters, including, for instance, a bimodal distribution in stage N2 unilateral spindles. N3 spindles and those in N2 coupled with SOs tended towards lower frequencies but still included a small proportion of spindles with frequencies ≥ 13 Hz, particularly in the young age group. Intriguingly, human intracranial recordings have shown narrow unimodal frequencies at each cortical location, against the backdrop of broader frequency distributions between different sites^18^. Additionally, an early simultaneous EEG and fMRI study on predefined fast and slow spindles identified several cortical differences in hemodynamic activity but found no differences in the thalamus, where spindle activity originates^96^. Collectively, these observations may suggest associations between spindling frequency and their capacity to propagate across multiple sites, possibly influenced by the structural or neurochemical architecture of these sites, which changes during ageing.

### Differences in spindle profiles in ageing

Known age-related differences in sleep architecture and spindle attributes were well captured in our sample. Specifically, we observed reduced sleep duration, a decreased proportion of stage N3 sleep, and lower spindle frequencies and absolute amplitudes. Unlike several previous studies^28,31,97,98^, we found no differences in spindle densities. This discrepancy may be explained by our inclusive approach, which combined all potential spindle types into a single density measure and normalization to the durations of specific sleep stages, along with our use of data-driven clustering rather than a priori selection of spindles based on specific attributes (e.g., slow or fast spindles).

As a novel aspect, we also explored post-spindle signal attributes. Despite spindles being optimal states for cellular-level plasticity, it has been noted that highly synchronous oscillatory events typically indicate periods of low information processing capacity^52^. Therefore, processes related to cortical replay may occur immediately after spindle events. We found that the global post-spindle signal dynamics differed between young and older age groups, with the older exhibiting a reduced 1/f slope of the power spectrum and decreased delta along with increased alpha and sigma frequencies. While non-linear signal complexity is typically decreased during sleep^57,91,99^ we found higher Katz fractal dimension in older adults, suggesting increased signal complexity. However, even though we extracted these values specifically after each spindle event, they were not distinguished from the broader context of the sleep stage. Therefore, the age differences in post-spindle signals possibly reflect global signal changes related to the transition between sleep stages and reduction of slow-wave sleep in ageing, rather than specific deviations in brain processing after each spindle event.

Data-driven clustering revealed novel insights into age-related changes in spindle activity. Specifically, we identified a proportional shift in older adults, characterized by a decline in bilateral stage N2 spindles and a comparative increase in N2 spindles that co-occurred with SOs. We speculate that this proportional shift was driven by a reduction in N2-SO^-^-bi clusters among older adults. Specifically increased proportion of N2-SO^+^ likely reflects the reduced deep sleep power, which makes it more likely for a given epoch to be scored as N2, rather than N3 stage in older adults^83^.

The reduction of N2-SO^-^-bi spindles is particularly intriguing, given the age-group-specific cluster attributes. Not only did older adults exhibit a smaller proportion of these spindles, but those that still belonged to this cluster also showed a significant frequency decline, with an almost complete separation between fast and slow spindles based on age group (Figure 7B). Additionally, a decline in frequency and power was observed among older adults across all clusters. These findings suggest that multiple mechanisms may contribute to the age-related decline of fast spindles^45^: deficiencies in spindle generation or propagation mechanisms, as well as a general slowing of all spindles, including those that otherwise display cluster-concurrent spatiotemporal profiles similar to the fast spindles found in young adults.

### Spindle associations with age-related memory decline and sleep consolidation effect

By combining several experimental procedures with assessments of subjective everyday memory complaints, we isolated age-related variance across multiple memory instruments. Parsing spindle heterogeneity into distinct spatiotemporal clusters revealed a cluster-specific association with the compound memory score and the proportion of N2-SO^-^-bi spindles in the younger age group. Notably, this is the same cluster that also showed age-related proportional shifts. After adjusting for age group differences, better memory in younger adults was associated with reduced proportion of N2-SO^-^-bi spindles, whereas older adults generally had lower proportion of those spindles. We speculate that this is reflects age-related shifts in the interdependent proportions of the N2-SO^+^ and N2-SO^-^-bi spindles, where the combination of the underlying SO phase angle and the ability to propagate across cortical areas may be relevant attributes for declarative memory.

Adding complexity to these findings, we observed associations between spindle power – irrespective of cluster membership – and the sleep-dependent memory effect, despite no age differences in this measure. The negative direction of this association (higher power related to reduced sleep gain), contrasts with previous findings. However, we calculated the sleep gain effect as the difference between sleep and awake conditions (in the absence of pre-learning baseline differences). Often, the sleep consolidation effect is calculated relative to pre-sleep learning performance^15^. Given the high correlations between performance after 12 hours of sleep and wakefulness (r ≈ 0.75), this difference measure may better capture sleep’s specific impact on memory consolidation. Here, this measure was positive, indicating reduced forgetting oversleep compared to wakefulness, consistent for both younger and older age groups. The modulation of the sleep consolidation effect by age remains controversial. Some studies report no sleep-related gains in older adults^3,35,59^, while others show similar gain in younger and older age groups^100–103^. Furthermore, potential publication bias for positive effects in young adults has been noted^59,60^. In conclusion, while our results affirm potential links between sleep spindles and memory performance, there is significant heterogeneity in these associations, and they seem vulnerable to subtle variations in memory assessment procedures.

The specific phase angle of SOs at the point of maximum spindle amplitude has gained feasibility as a potential predictor of age-related memory recline ^43,44,104^. Overall, we replicated previous findings, in which the spindles preferably occur in the up-phase closer to the positive peak in younger adults, whereas this distribution was more spread and shifted towards the negative peak in older adults. However, our clustering results revealed some paradoxes on spindle SO-coupling and aging.

Specifically, deviant coupling and its related memory decline have only been demonstrated for fast spindles^43^. However, concurrent spindle and SO events are relatively rare. In our results, stage N3 spindles (with increased SO activity) and stage N2 spindles that coincided with SOs had frequency distributions skewed towards the typical range of slow spindles (< 13 Hz). Additionally, aging significantly affected frequencies across all clusters, resulting in a notably reduced high-frequency peak in the probability density functions compared to the younger age group. These factors diminish the concurrence of fast spindles and SOs in aging on top of an already small baseline. In this vein, findings from a large community sample showed that only about half of individuals over 50 years of age had above-chance concurrence of SOs and sleep spindles^40^. Thus, despite the concept of deviant SO and fast spindle phase coupling in aging being well-documented and replicated here, the rarity of these events in older adults suggests caution in interpreting this measure mechanistically. In the context of recent large-scale findings indicating that many interrelated sleep parameters similarly predict cognitive decline^40^, it’s plausible that deviant SO-spindle coupling is an epiphenomenon reflecting a breakdown of global network dynamics, potentially driven by structural brain changes or functional alterations in deep sleep during aging. Nonetheless, as SO-spindle coupling has been associated with increased beta-amyloid burden and predicted cognitive decline at least two years in advance^105^, it may still hold promise as a biomarker to detect accelerated aging and pathology risk in midlife, when the deterioration of fast spindles is only beginning to emerge.

### Spindle attributes are intertwined with global sleep dynamics

Our study elucidates the intricate relationship between spindle attributes and global sleep dynamics, particularly as they change with aging. The calculation of absolute spindle power is inherently influenced by the 1/f power spectrum^106^, while relative spindle power tends to decrease as low frequencies in deep sleep start to dominate. The global power spectrum and the proportion of slow wave sleep shifts significantly with aging, complicating the interpretation of spindle power measurements. Additionally, the power fluctuations across the night and spindling frequency also seem to align with sleep stage transitions from light N2 towards periods with more SO until reaching stage N3, which is dominated by SOs. While some spindles exhibit unique propagation across the brain, possibly indicating specialized functions, this too appears influenced by SO dynamics that facilitate cortical travel under specific conditions^21,107^. Taken together, these insights suggest that many previously noted spindle-specific attributes may be more accurately seen as manifestations of broader changes in sleep architecture with aging.

### Limitations

Our study has several limitations. First, although we focused our analysis on the electrodes most used in clinical polysomnography, the absence of frontal and parietal electrodes may limit our ability to distinguish between slow and fast spindles. Second, the spindle heterogeneity is likely even greater than presented here, as EEG cannot detect local spindles captured intracranially or propagation through core thalamocortical projections, which can be observed with magnetoencephalography^50,51^. Third, the final clustering solution heavily depends on the initial features and the criteria used to determine the number of clusters. By including interrelated temporal features, our analysis might be biased toward sleep stage information. Therefore, it is unlikely that the resulting clusters capture spindles that are truly mechanistically and/or functionally distinct. However, the spatiotemporal clustering revealed in our analysis underscores that spindle attributes are not independent of the global spectral context. Lastly, our sample size lacks sufficient statistical power to reliably detect small memory associations, particularly across multiple features. At the same time, testing multiple interrelated measures hampers the interpretation of significance due to reduced control over type I error. This is a broader issue in the field, especially in studies comparing several spindle features in small patient groups. A recent large-scale study suggests that multiple interrelated spindle features are related to cognitive function in aging, but also to other sleep variables^40^. This challenges the proposition that sleep spindles could serve as more specific biomarkers of early pathological aging than other sleep parameters.

## Conclusion

In conclusion, our data-driven clustering approach sought to identify distinct spindle types with functional significance in relation to aging and memory decline. We identified several spatiotemporal clusters, highlighting notable age-related differences, such as a reduced proportion of stage N2 bilateral spindles and an increased proportion of N2 SO-concurrent spindles, along with a general decline in frequency and power. However, our clustering solution represents just one possible way of categorizing spindles, influenced by the initial feature set and the final choice of cluster numbers. Whereas none of the clusters emerged as a definitive predictor of memory decline in older age, the observed age differences and memory associations in the younger age group highlight the intertwined dynamics between spindle propagation, SO concurrence, and frequency shifts in ageing. Moreover, our findings emphasize the importance of considering spindle activity within the broader context of sleep architecture, recognizing their role within global network dynamics and overall sleep patterns.

### Data and code availability

Individual-level data availability is restricted as participants have not consented to publicly share their data. The data may be available upon reasonable request, given appropriate ethical, data protection, and data-sharing agreements. Requests can be submitted to the principal investigator (Prof. Anders Fjell, University of Oslo). All analysis scripts are available at the dedicated project at Open Science Framework (https://osf.io/k6wg3/).

## Acknowledgments

This project was funded by European Research Council to A.M.F under grant agreements no. 725025 and the Norwegian Research Council no. 325878. B.R was funded by Swiss National Science Foundation (SNSF) (10.002.483).

