## Supplementary Materials for "Differential expression of spatiotemporal sleep spindle clusters in ageing"


Liisa Raud[1], Martijn J. B. Smits[1,2], Markus H. Sneve[1], Hedda T. Ness[1], Line Folvik[1], Björn Rasch[3], Anders M. Fjell[1,4]

1 Center for Lifespan Changes in Brain and Cognition, Department of Psychology, University of Oslo, 0373 Oslo, Norway

2 Clinical Neuroscience Laboratory, Department of Psychology, Norwegian University of Science and Technology, 7491 Trondheim, Norway

3 Department of Psychology, Division of Cognitive Biopsychology and Methods, University of Fribourg, Switzerland

4 Computational Radiology and Artificial Intelligence, Department of Radiology and Nuclear Medicine, Oslo University Hospital, 0372 Oslo, Norway

**Supplementary materials 1: Pruning of the feature matrix**

(1) Correlation-based pruning

Pearson correlation matrix was calculated across all continuous features and, of the pairs with $r > 0.70$, only one of the features were retained. As a result, five features were discarded: amplitude and root mean square (triangular correlations with absolute power), number of oscillations (correlated with duration), Petrosian fractal dimension (correlated with permutation entropy), and post-spindle spectral density intercept (correlated with the exponent).

(2) Post-hoc pruning

Second pruning of the feature matrix was done after initial results were obtained. First, it was noted that the categorical feature 'cycle' and continuous variable 'time relative to sleep onset' were highly discriminative between clusters. The clustering was thus repeated after omitting 'cycle' and retaining the continuous timing variable, to obtain results without the partial redundancy between these two features. Lastly, it appeared that clustering results were biased by the stage information, likely due to the inclusion of numerous inter-related post-spindle period variables. Therefore, the clustering was repeated after discarding all post-spindle period variables. We chose to report this analysis as the main results for several reasons: (1) the stage information was captured well also in the results without post-spindle variables, and (2) as the variables originated from the signal *after* the spindle events, there was ambiguity whether we were clustering spindles or merely differences in sleep stages.

**Supplementary materials 2: Source Memory Effects Separately for Sleep and Awake Conditions**

| Memory measure | Level of specificity | Young | | Old | | r sleep-awake |
|---|---|---|---|---|---|---|
| | | Sleep | Awake | Sleep | Awake | |
| Baseline | item | 76 (17) | 71 (21) | 58 (21) | 55 (20) | 0.73 |
| 12 hours source recall* | category | 72 (21) | 65 (22) | 53 (27) | 49 (26) | 0.72 |
| 12 hours recall relative to baseline performance | item | 88 (10) | 81 (14) | 73 (16) | 65 (19) | 0.76 |
| 6 days source recall | item | 54 (24) | 50 (25) | 34 (18) | 28 (17) | 0.76 |

Table S1. Memory measures for the source memory experiment, separately for awake and sleep condition. The values are means (standard deviations in brackets) per group and condition. Note that these are all original measures, without imputations. The number of missing values per cells varies between 0-3. The level of specificity indicates whether correct recall is considered at the item-level (i.e. correct recollection of the specific face or place) or category level (i.e. face or place). *indicates the measure which formed the basis for calculating the sleep gain effect in the main analysis, subtracting the sleep from the awake performance for each individual.

**Baseline** indicates the percentage of correct items directly after encoding (AFC-1), before the 12 hour sleep/awake period. **12 hours source recall** indicates the percentage of correct associations after the 12 hour sleep/awake period. **The 12 hours recall relative to baseline performance** indicates correct associations at baseline and 12 hours (AFC-1-hit & AFC-2-hit) as a percentage of the baseline (AFC-1) performance, and indicates the percentage of remembered items from initially learned items. **6 days source recall** indicates the percentage of correct associations that were correct at all three test times (AFC1-hit & AFC-2-hit & AFC-3-hit): baseline, 12-hours, and 6 days.

For each of the variables, we ran linear regression models, predicting memory from the age group (young vs old), condition (sleep vs awake), and interaction. In each case (except for the baseline, as reported in the main text), we found significant main effects (young > old, sleep > gain), but no interactions, indicating stability of findings across different ways of calculating the memory performance.

**Supplementary materials 3: Sleep architecture**

*Methods*

Main indicators of sleep architecture were calculated per age group: sleep period time (from sleep onset to awakening), total sleep time, wake after sleep onset, and the percentages of time spent in each sleep stage (N1, N2, N3, REM). In addition, the average count of spindles per person was calculated, as well as overall spindle densities, normalized to the summed duration of NREM sleep stages N2 and N3. These values were compared between young and old age groups using independent t-tests. One participant was discarded from the global sleep architecture analysis due to bad signal after 4.5 hours or sleep.

*Results*

| **Variable** | **Old** | **Young** | **t (df)** | **Cohen's d** | **p-value** |
|---|---|---|---|---|---|
| Sleep period time (h) | 5.70 (0.73) | 6.16 (0.56) | -2.99 (58) | -0.720 | 0.004* |
| Total sleep time (h) | 5.10 (0.74) | 5.70 (0.48) | -4.07 (52) | -0.99 | <0.001* |
| Wake after sleep onset (min) | 36 (25) | 27 (20) | 1.64 (62) | 0.390 | 0.107 |
| N1 proportion (%) | 8.4 (4.1) | 5.2 (2.7) | 3.78 (53) | 0.920 | 0.000* |
| N2 proportion (%) | 55 (10) | 48 (5) | 3.48 (47) | 0.870 | 0.001* |
| N3 proportion (%) | 17 (9) | 25 (6) | -4.07 (52) | -0.99 | 0.000* |
| REM proportion (%) | 19 (7) | 21 (5) | -1.52 (52) | -0.370 | 0.134 |
| Spindle count (#) | 1,223 (511) | 1,340 (440) | -1.06 (65) | -0.250 | 0.293 |
| Spindle density (# per minute of NREM sleep) | 5.58 (1.89) | 5.33 (1.65) | 0.59 (64) | 0.140 | 0.558 |

Table S2.Sleep variables and statistical comparisons between young (n=43) and old (n=34) age groups. The values for young and old are mean values, with standard deviations in the brackets. *marks significant difference between young and old at an alpha level of 0.05.

**Supplementary materials 4: Alternative clustering pipelines**

*Group normalization*. For comparable distances between variables in the clustering algorithm, all variables must be normalized between zero and one. In our original analysis, the variables were normalized within each person. The advantage of this is that the final cluster centroids are less dependent on the original sample. That is, each new individual stands as an independent contribution, so that the final solution can be more generalizable across different samples. However, the limitation is that inter-individual variability is reduced, which may obscure the memory predictions. We therefore repeated the analysis after normalizing continuous variables at the group level.

With this normalization scheme, the variable significances were similar with the original solution, with stage and relative power having the strongest influence and k = 4 providing the best fit. In addition, the cluster profiles were almost identical, with the exception that the N2-bilateral cluster also included some spindles that co-occurred with the SOs, whereas the N2-$SO^+$ cluster consisted of only unilateral spindles. The memory results were similar to the original solution, in which the age group and proportion interaction was significant for the N2-$SO^-$-bi cluster and the power correlated negatively with the sleep gain effect in each cluster.

*Inclusion of post-spindle signal features.* Initially, a set of features from the post-spindle signal of two seconds were extracted, based on the reasoning that spindles may serve as a gating events for following cortical consolidation processes. These features were later discarded, due to the perceived bias towards sleep stage classification. When including these features, sleep stage was still the variable with the largest significance, followed by other features capturing the power spectrum of the post-spindle signal (delta power, 1/f exponent, sigma power, alpha power, Katz fractal dimension, and theta power). Here, both k = 2 and k = 3 provided better fits than higher k's. With k = 3, clusters separated into stage N3 spindles, N2-SO-concurrent spindles, and a single cluster including all N2 spindles that did not co-occur with SOs. No age-related memory associations (PC1) were significant with this solution. The sleep gain associations reflected the original results, with negative associations between spindle power and the sleep gain effect in each cluster.

**Supplementary materials 5: SO-sigma coupling**

*Methods*

The spindle-SO coupling was first calculated for all SOs using the YASA toolbox[1]. Shortly, the broadband data was first filtered in the SO-band (0.3-1.5 hz) and the instantaneous phase angle was extracted after the Hilbert transform of the -1/+1 second segments around the negative peaks. Then, the same data was filtered in the broad sigma frequency band (9-16 Hz) and for each SO, the phase angle at the time of the maximum sigma amplitude was extracted. Next, for each spindle that had previously determined to concur with the SO, we retained the SO with a negative peak closest to the spindle peak. The mean phase angle was calculated per participant and cluster using the *circular*[2] package in R and is expressed in radians with -pi/pi corresponding to the SO negative peak, 0 to positive peak, and positive and negative values indicating down- and up-phase, respectively.

For statistical testing of the distribution of the SO phase angles, the full SO cycle in radians was divided into 12 bins, and for each participant and cluster, the mean number of spindles per bin was calculated. For statistical testing, we shuffled the temporal order of the bins for each participant 1000 times, averaged over the permutation iteration for each bin, tested each bin against this permuted surrogate value using dependent t-tests, and applied false discovery rate p-value correction over the 12 bins[3]. This procedure was done separately for the old and young age groups. Further, the age differences in each cluster were tested by comparing the mean phase angles with Watson-Williams tests, which is a t-test alternative for circular data, using the *circular*[2] R package. Additionally, circular correlation between the phase angles in each of the two clusters was calculated using the *CircStats*[4] package in R. Lastly, circular-linear regression coefficients were calculated for the associations between the mean phase angle and age-related memory performance, as well as between the mean phase angle and sleep-related memory gain, using the *directional*[5] R-package.

*Results*

In line with previous literature, the N2-SO+ spindles preferably occurred just before and at the positive peak in young adults, with lower accounts on the down-phase (see figure below). The N3 spindles in the younger age group had a more uniform distribution on the up-phase, but still had lower occurrences on the down-phase than chance level. The older adults had more varied distributions, with spindles still occurring rather on the up phase, but closer to the negative peak. These relationships were also reflected by the mean phase angles across age groups (Figure 9B). Whereas these occurred at the up-phase for both clusters and age groups, the spindles in younger adults were closer to the positive peak than in the older adults both in the N2-SO+ cluster (mean phase angle old: -2.58 rad, young: -1.21 rad; Watson-Williams circular test F = 25.55, df = 1, 75, $p<0.001$) and the N3 cluster (old: -2.76 rad, young: -1.55, F = 10.49, df = 1, 75, p = 0.002). Note that the mean phase angles correlated between the N2-$SO^+$ and N3 clusters (circular r = 0.53, $p < 0.001$), and this relationship was preserved in both age groups when tested separately.

We tested the relationship between the SO phase angle and the memory performance using circular-linear regressions (panel C). We found significant relationship between the phase angle of the N2-$SO^+$ cluster and the age-related memory performance captured by PC1 ($R^2$ = 0.07, p = 0.006), with participants with the phase angles closest to the positive peak on the up-phase showing better memory performance (panel C; upper left). However, this relationship was likely driven by the age differences in both variables, as it was not significant if tested separately in the age groups (both $R^2$'s $< 0.01$, p's $>= 0.827$). There were no other significant memory associations neither with the PC1 nor with the sleep gain (all R's $<= 0.03$, p's $>= 0.111$). Note, however, that when tested separately in each age group, there was a relationship between the phase angle of the SOs coupled with the N3 spindles and the PC1 in the older adults ($R^2$ = 0.11 , p = 0.032), in which older adults with SO phase angles closest to the negative peak had better memory performance (panel C, upper right).

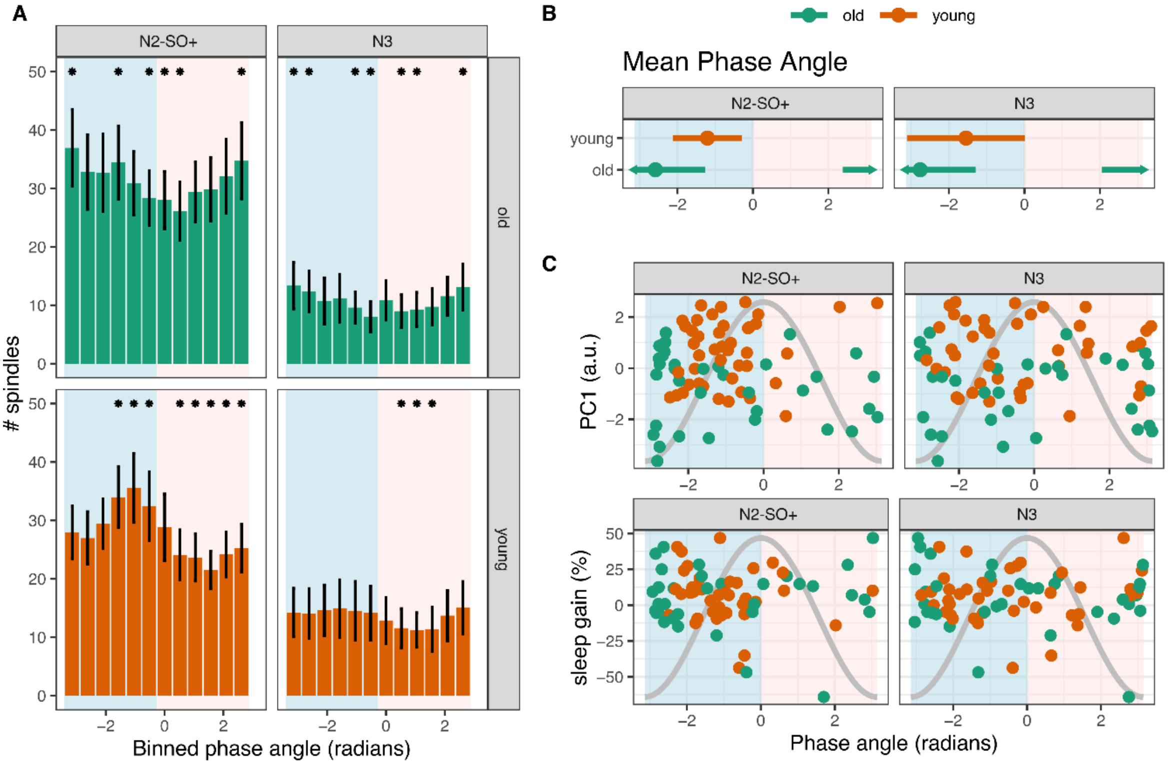


Figure S1. (A) Mean number of spindles per SO phase angle bin, divided into 12 bins. The error bars indicate 95% confidence intervals. *marks significant difference when tested against within-participant permuted surrogate data. (B) Mean phase angles, with the error bars representing standard deviations. The arrows on the error bars in the older group indicate continuation over the negative peak at +/-pi. (C) Relationship between memory associations and the SO phase angle. In all subfigures, blue background shading indicates the SO up-phase and pink shading indicates the down phase.

**Supplementary materials 6: Spindle power and sleep gain associations by cluster**

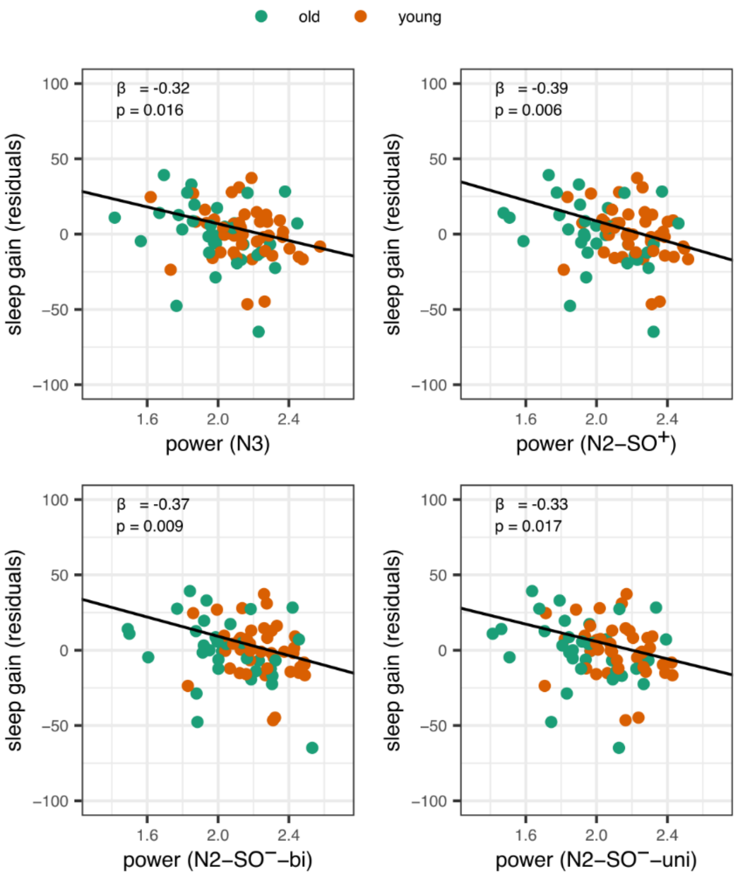


Figure S2. Associations between sleep gain and spindle power in each cluster, regardless of age group. The sleep gain memory scores are residualized for sex and age group, but the regression lines and text annotations represent the intercept, slope, beta, and p-values from the original models.
